# A ChIP-exo screen of 887 PCRP transcription factor antibodies in human cells

**DOI:** 10.1101/2020.06.08.140046

**Authors:** William K. M. Lai, Luca Mariani, Gerson Rothschild, Edwin R. Smith, Bryan J. Venters, Thomas R. Blanda, Prashant K. Kuntala, Kylie Bocklund, Joshua Mairose, Sarah N Dweikat, Katelyn Mistretta, Matthew J. Rossi, Daniela James, James T. Anderson, Sabrina K. Phanor, Wanwei Zhang, Zibo Zhao, Avani P. Shah, Katherine Novitzky, Eileen McAnarney, Michael-C. Keogh, Ali Shilatifard, Uttiya Basu, Martha L. Bulyk, B. Franklin Pugh

**Author notes:** **Contact Information:** William KM Lai < >, Luca Mariani < >, Gerson Rothschild < >, Edwin R. Smith < >, Bryan J. Venters < >, Thomas R Blanda < >, Prashant K Kuntala < >, Kylie Bocklund < >, Joshua Mairose < >, Sarah N Dweikat < >, Katelyn Mistretta < >, Matthew J. Rossi < >, Daniela James < >, James T. Anderson < >, Sabrina K. Phanor < >, Wanwei Zhang < >, Zibo Zhao < >, Avani P. Shah < >, Katherine Novitzky < >, Eileen McAnarney < >, Michael-C. Keogh < >, Ali Shilatifard < >, Uttiya Basu < >, Martha L. Bulyk < >.

## Abstract

Antibodies offer a powerful means to interrogate specific proteins in a complex milieu. However, antibody availability and reliability are problematic and epitope tagging can be impractical in many cases. In an effort to improve this situation, the Protein Capture Reagents Program (PCRP) generated over a thousand renewable monoclonal antibodies (mAbs) against human-presumptive chromatin proteins. However, these reagents have not been widely field-tested. We therefore performed a screen to test their ability to enrich genomic regions via chromatin immunoprecipitation (ChIP) and a variety of orthogonal assays. 887 unique antibodies against 681 unique human transcription factors (TFs), were assayed by ultra-high resolution ChIP-exo/seq, primarily in a single pass in one cell type (K562). Deep systematic analyses of the resulting ∼1,200 ChIP-exo datasets can be found at www.PCRPvalidation.org. Subsets of PCRP mAbs were further tested in ChIP-seq, CUT&RUN, STORM super-resolution microscopy, immunoblots, and protein binding microarray (PBM) experiments. About 5% of the tested antibodies displayed target (i.e., cognate antigen) enrichment across at least one assay and are strong candidates for additional validation. An additional 34% produced ChIP-exo data that was distinct from background and thus warrant further testing. The remaining 61% were not substantially different from background, and likely require consideration of a much broader survey of cell types and/or assay optimizations. We demonstrate and discuss the metrics and challenges to antibody validation in chromatin-based assays.

## Introduction

Antibodies are a critical component of a wide variety of biochemical assays. They serve as protein-specific affinity-capture and detection reagents, useful *in vivo* and *in vitro*. Example assays include chromatin immunoprecipitation (ChIP) of protein-DNA interactions, immunofluorescence, immunoblotting, ELISA, purification of cells and proteins, protein binding microarray (PBM) experiments, and targeted *in vivo* delivery of effector molecules (Chames et al. 2009; Park 2009; Siggers et al. 2011a; Mahmood and Yang 2012; Engelen et al. 2015; Lin et al. 2015). One advantage of target-specific antibodies is their ability to recognize proteins without the need for an engineered affinity tag. The human proteome contains tens of thousands of distinct proteins, each requiring a different antibody for specific detection. The usage of a variety of antibodies to diverse targets has been a critical component of NIH-funded consortium projects such as the ENCODE and Roadmap Epigenomics Mapping (Consortium 2012; Roadmap Epigenomics et al. 2015). However, broad profiling of the genomic targets of human sequence-specific transcription factors (ssTFs) has been limited by the availability of ‘ChIP-grade’ antibodies.

Overall, there has been an acute lack of antibodies that effectively distinguish the many thousands of different chromatin proteins. Consistency in reagent production and performance have been particularly problematic (Egelhofer et al. 2011; Baker 2015; Shah et al. 2018). Polyclonal antibodies, being a mixed product of many antibody genes, have the advantage of potentially recognizing multiple epitopes on a protein, thereby producing robust target detection (Hanly et al. 1995). However, their production is finite, and can be variable across immunized animals, but also vary within individuals by different bleed dates and affinity purifications. These factors and more hamper reproducibility (Reardon 2016).

The NIH Protein Capture Reagent Program (PCRP) was initiated through the NIH Common Fund with the stated goal of testing the feasibility of producing low-cost, renewable, and reliable protein affinity reagents in a manner that can be scaled ultimately to the entire human proteome (PA-16-287) (https://proteincapture.org/) (Blackshaw et al. 2016). With an initial focus on putative ssTFs, this endeavor reported the production of 1,406 mouse monoclonal antibodies (mAbs) against 737 chromatin protein targets (Venkataraman et al. 2018). This included two parallel production approaches: mouse hybridomas that release mAb into growth medium supernatant, and recombinant antibodies produced in *E. coli*. The advantages of these two approaches over polyclonal antibodies are, in principle, a renewable and consistent supply of homogeneous preparations produced from a single set of genes that recognize a single epitope (Kohler and Milstein 1975; Winter et al. 1994; Liu 2014). To accommodate the potential shortcoming of a hybridoma recognizing a single non-viable epitope, the NIH PCRP made an effort to generate at least two independent clones for each target, although this does not guarantee two different epitopes, as in cases where there are immunodominant regions.

Antibody validation is required to generate confidence in their utility (Baker 2015; Marx 2019). Validation exists at many levels ranging from whether an antibody specifically recognizes its intended target to the exclusion of all others, to whether it consistently performs successfully in a particular assay (Landt et al. 2012; Wardle and Tan 2015; Uhlen et al. 2016; Edfors et al. 2018; Sikorski et al. 2018). Each publicly available PCRP-generated antibody was previously validated for its target recognition by *in vitro* human protein (HuProt) microarray screening (Venkataraman et al. 2018). These arrays contain approximately equivalent amounts of antigen, which differs from the wide expression range in natural sources. They also may differ in epitope accessibility compared to complexed or crosslinked targets in ChIP assays. Thus, additional assay-specific validation is necessary. The further capability of PCRP antibodies has been described for a limited set in a number of approaches, including immunoblotting, immunoprecipitation, immunohistochemistry, and ChIP-seq, with assay-dependent success observed (Venkataraman et al. 2018). Previous work has reported that 46 of 305 mAbs against 36 of 176 targets passed ENCODE ChIP-seq standards (Venkataraman et al. 2018), although detailed supporting evidence is not publicly available. “Browser shots” of selected loci from that study are available for ∼40 datasets against 31 targets. However, these should be considered preliminary, since locus-specific examples lack statistical power and unbiased selection. Additionally, chromatin fragmentation and extraction may generate localized variation in yield that varies from prep to prep (active promoters, enhancers, etc.). Numerous replicates (target and control) are often needed to ensure against false positives at selected individual loci due to sampling variation and multiple hypothesis testing. As broader community use of the PCRP-generated antibodies will likely benefit from a wider survey, we conducted additional tests of these reagents. To our knowledge there has been no systematic large-scale field assessment of antibodies in ChIP. We report on progress and challenges in comprehensively assaying ∼1,400 PCRP mAbs. As this represents a first-pass assessment, most experiments include only a single replicate, using enrichment of specific genomic features as preliminary evidence of success. We performed replicates on some samples that displayed enrichment (e.g., expected motif enrichment) as well as a subset of samples which displayed no initial enrichment to examine our true negative rate.

Since evaluating each of ∼1,400 mAbs in a wide variety of assays was not practical, we opted for broad coverage by ChIP-exo, which we have developed into a high-throughput and ultra-high resolution alternative to ChIP-seq (Rossi et al. 2018). ChIP-exo allows genome-wide detection of chromatin interactions at near-bp resolution, which also increases the confidence of peak calling. We further tested a smaller subset of mAbs in other assays (ChIP-seq, CUT&RUN, super-resolution cellular microscopy (STORM), immunoblots, and protein binding microarrays). These additional tests were not intended to be comprehensive, but rather to evaluate the challenges and practicality of systematic antibody validation. Overall, we tested 946 unique mAb clones (887 in ChIP-exo and 59 in other assays), of which 642 targeted putative ssTFs, which allowed computational comparison through the enrichment of their cognate motifs (if one exists). The antibodies and assays were chosen to cover a wide range of end-user applications, with specific ssTFs chosen in part based on the scientific interests of the investigators and a set of objective criteria. With a deep dive on a single assay (ChIP-exo), we explored end-user practical issues related to antibody sourcing, reproducibility and validation metrics, and specificity for cell types and states.

## Results

### Screening PCRP mAb by ChIP-exo

We used the massively parallel ChIP-exo version of ChIP-seq to screen PCRP Abs in 96-well plate format (48 at a time) for their ability to recognize their putative protein targets in a chromatinized, cellular context (Rhee and Pugh 2012; Rossi et al. 2018). Briefly, proteins were formaldehyde crosslinked to DNA and each other within cells. Chromatin was then isolated, fragmented, and immunoprecipitated. While on the beads, the fragmented DNA was trimmed with a strand-specific 5’-3’ exonuclease up to the point of crosslinking (i.e., protection), which was then mapped by DNA sequencing. For many proteins, this provides single-bp resolution in genome-wide detection (Rhee and Pugh 2011). Since ChIP-exo is a higher resolution derivative of ChIP-seq, ChIP-exo is expected in principle to detect any real binding events that ChIP-seq detects, and likely more due to its higher dynamic range.

Technical reproducibility of ChIP-exo with PCRP mAbs was evaluated with 43 independent replicates performed on the sequence-specific TF USF1. This replicate served as a positive control in 43 cohorts, each having 46 mAb assayed on different days: as such no USF1 replicates were excluded in this analysis. We prioritized mAb evaluation in K562 (human bone marrow lymphoblast), but also tested a subset in MCF7 (human mammary gland epithelial), HepG2 (human liver, epithelial-like), and donated human tissues (liver, kidney, placenta, and breast). From this and IgG (or no antibody) negative controls, we defined with USF1 a set of ∼164,000 E-box motif instances associated with a significant (q<0.01) peak-pair in at least one replicate ChIP-exo experiment (Albert et al. 2008). This reflects a very relaxed criterion, with a high level of expected false positives, for the purposes of evaluating the gradient from true binding through nonspecific background at genomic E-boxes. The latter is expected where a peak location occurred in only a small fraction of datasets. When examined at greater stringency regarding the number of replicates in which the same peak was found, a higher average occupancy and more robust patterning was observed. Of the USF1 datasets, 43 of the 45 (>95%), produced a USF1-specific ChIP-exo pattern around E-boxes (*Supplementary Fig. 1A*, vertical blue and red stripes in the heat maps and single-bp peaks in the composite plots). A Pearson-pairwise correlation was calculated for the occupancy of binding at putative USF1-bound E-boxes across all USF1 and negative control IgG (or no antibody) ChIP datasets (*Supplementary Fig. 1B*). A strong correlation among USF1 data sets was observed, which reflects a high level of reproducibility and indicated that ChIP-exo was suitable for screening and evaluating PCRP mAbs in ChIP.

We initiated our ChIP mAb survey by first considering the practicality of producing a large number of antibody preparations from *E. coli* or mouse hybridomas. Purification from *E. coli* included transformation of expressing plasmids, cell growth, recombinant protein induction, and purification. After multiple attempts, we determined that high throughput parallelized recombinant immunoreagent production was not practical within the scope of this project, due to a need to optimize the protocol in our hands. We therefore opted to pursue commercially available hybridoma-based mAbs.

We first examined vendor source. We tested NRF1, USF1, and YY1 PCRP mAb from DSHB and CDI. The former was supplied as hybridoma culture supernatants (10-80 ug/ml antibody), and the latter as concentrates (*Supplementary Fig. 2*). Each preparation (as supplied) was pre-loaded onto protein A/G magnetic beads. In general, we found that while mAbs from both sources (assayed at the same reported mAb amounts; 3 ug) specifically detect NRF1 and USF1, DSHB-derived hybridoma culture supernatants detected more binding events at cognate motifs compared to CDI concentrates. We therefore sourced from DSHB for the remainder of this study. Nevertheless, mAbs from CDI may be improved through further optimizations.

We assayed all 887 available hybridoma supernatants containing mAbs to 681 non-redundant targets. Testing was initially performed in K562 (1,009 datasets), although a subset of hybridoma supernatants were tested in MCF7 (134 datasets) and HepG2 (96 datasets) cells, based on reported target mRNA expression levels. If there was no substantial difference in ssTF expression level, or the ssTF was not measurably expressed in any of these three lines, testing defaulted to K562 due to practical considerations including the ability to grow this line at scale (liquid culture) (*Supplementary Fig. 3A*). 245 unique hybridoma clones were assayed in replicate at least twice in the same or different cell types, resulting in 1,261 datasets (*Supplementary Fig. 3B*). Of these 245 hybridoma clones, 36 (14.7%) showed enrichment of the same class of genomic features. 102 (41.6%) of the mAb clones produced no enrichment of genomic features in both replicates, and 107 (43.7%) produced enrichment in one sample but not in the other (*Supplementary Table 1*). The latter may be due to being at the limits of detection. We next set out to characterize certain mAb in more depth.

As exemplified by NRF1, USF1, YY1, and an IgG negative control (**Figure 1A**), the finding that ChIP-exo peaks were enriched at a very precise distance from cognate motifs provided strong support for specificity in target detection. We also compared different hybridoma clones against the same target, as exemplified by heat shock factor 1 (HSF1). Hybridoma clones potentially target different epitopes, although immunodominance may yield independent clones to the same epitope. Both HSF1 mAbs gave nearly identical ChIP-exo read patterns around the same set of features (heat shock elements, **Figure 1B**, clones 1A10 and 1A8), thereby demonstrating reproducibility of ChIP-exo profiles across independent PCRP mAbs. This is particularly important where validation criteria by motif enrichment are not applicable. However, two other HSF1 mAb clones failed (**Figure 1B**, clones 1C1 and 1D11), indicating that independent PCRP clones can have different capabilities in ChIP. Therefore, if one mAb clone fails, it may be productive to check others.

**Figure 1.**
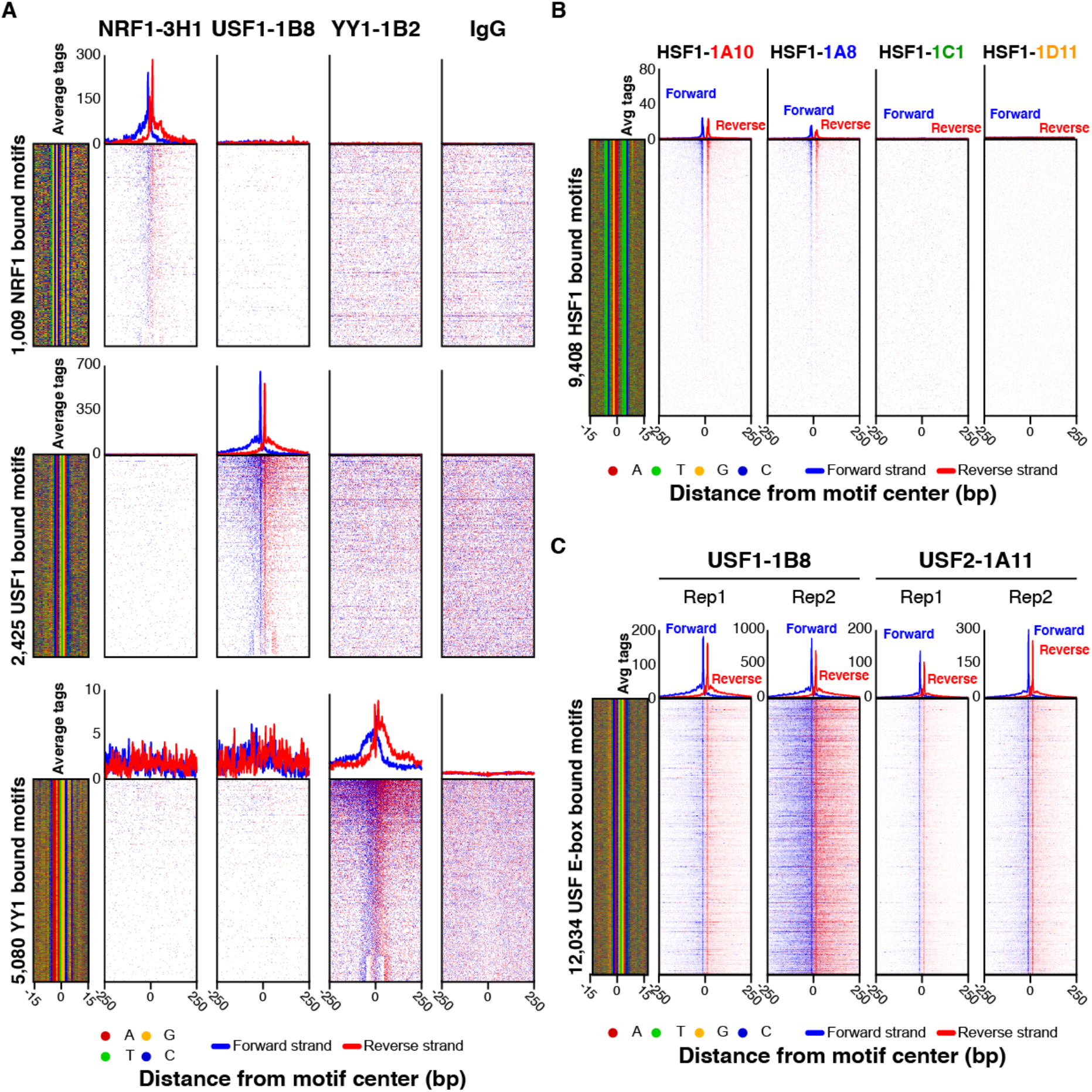
(**A**) Comparison of ChIP-exo data at cognate vs non-cognate motifs. ChIP-exo heatmap, composite, and DNA-sequence 4-color plots were generated for NRF1, USF1, YY1, and IgG ChIP-exo datasets against the complete matrix of bound motifs from *Supplementary Fig. 2*. The 5’ end of aligned sequence reads for each set of experiments were plotted relative to distance from cognate motif for each indicated target. Reads are strand-separated (blue = motif strand, red = opposite strand). Rows are linked across samples and sorted based on their combined average rank-order in a 100 bp bin around each motif midpoint. High levels of background result in a more uniform distribution of reads across the window (as seen with the IgG control). (**B**) Independent hybridoma clones and target interaction partners as antibody validation criteria. ChIP-exo heatmap, composite, and DNA-sequence 4-color plots are shown for the indicated number and type of bound motifs for (**A**,**B**) the indicated antibody hybridoma clones or (**C**) interaction partners, tested in K562 cells. The 5’ end of aligned sequence reads for each set of experiments were plotted against distance from cognate motif, present in the union of all called peaks between the datasets for each indicated target. Reads are strand-separated (blue = motif strand, red = opposite strand). Rows are linked across samples and sorted based on their combined average rank-order in a 100 bp bin around each motif midpoint.

Targets that interact with each other or with the same sites may also provide a useful validation criterion for determining enrichment specificity. For example, in the case of USF1 and USF2 interaction partners and homologs (Rada-Iglesias et al. 2008), the USF1-1B8 and USF2-1A11 mAbs detected binding at the same sites (**Figure 1C**). However, in this particular case we cannot exclude cross-reactivity of the mAb with the two homologous USF1/2 proteins (always a potential concern with target-specific antibodies). Additional validation criteria may include comparisons to public-domain ChIP-seq datasets that use different antibodies (*e.g.,* ENCODE, as in *Supplementary Fig. 4*).

### Assessment by ChIP-seq

Since ChIP-seq is a widely used assay (and related to ChIP-exo), we performed ChIP-seq (in HCT116 cells) with 137 PCRP hybridomas corresponding to 70 targets associated with chromatin binding, modification, enhancer function, and/or transcriptional elongation. We found 19 (14%) produced significantly enriched peaks (see Methods) (*Supplementary Table 2*). However, these single-replicate datasets were not checked for enrichment of specific classes of genomic features. Stringent validation for antibody-specificity involves knocking down a target, then observing a reduction of assayed signal relative to a mock knockdown (Wardle and Tan 2015; Uhlen et al. 2016; Edfors et al. 2018). We examined the feasibility of this starting with one target, NRF1. NRF1 expression was knocked down in HCT116 cells by RNAi. NRF1 peaks were concomitantly diminished with two different specific oligos, but not by an untargeted oligo (**Figure 2A**), thereby demonstrating specificity of the PCRP mAbs 3D4 and 3H1for this target. We further confirmed specific knockdown of NRF1 by immunoblot (**Figure 2B**). Due to the relatively high cost and current limitations in knock-down technologies, we found it was not practical within the scope of this project to conduct systematic knockdowns across the PCRP mAb collection. Furthermore, knockdown validation may not provide the level of validation stringency in ChIP that it does for immunoblots. Knockdown of proteins can cause widespread indirect effects on the binding of other protein-complexes which could in turn skew the ChIP-signal in aberrant ways (Trescher and Leser 2019).

**Figure 2.**
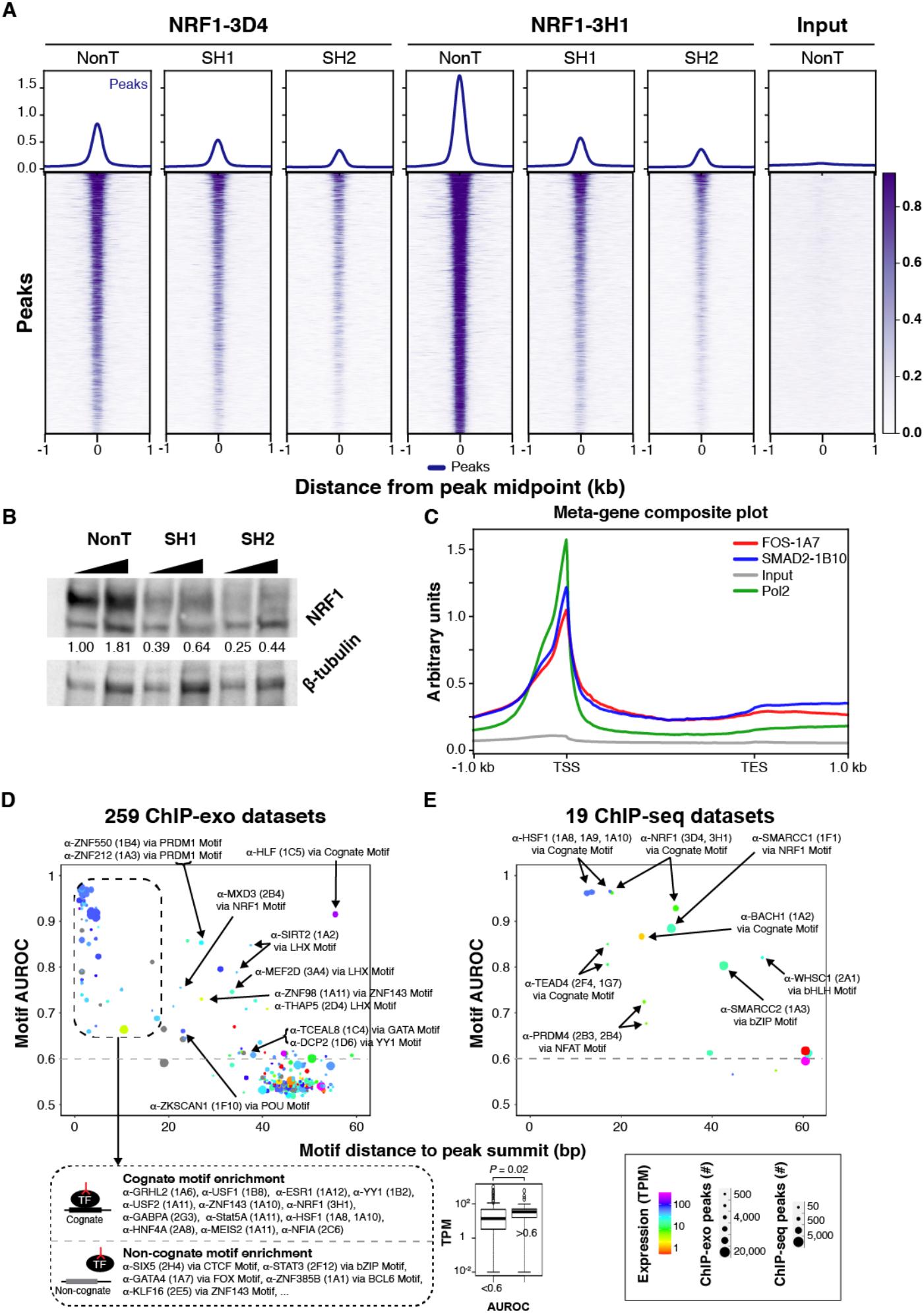
(**A**) Heatmaps and composite plots displaying the global loss of NRF1-3D4 and NRF1-3H1 ChIP-seq signal after NRF1 RNAi. ChIP-seq in HCT116 cells treated with non-targeting (sh Control) or two different NRF1-directed shRNAs (shRNA 1 and shRNA 2). Rows are linked across samples and sorted in descending order by mean score per region. (**B**) Western blot analysis of NRF1 knockdown by two different shRNAs (SH1 and SH2) or a non-targeting shRNA (NonT). HCT116 cells were infected with the indicated shRNAs and selected with puromycin (2 µg/ml). Total cell extracts were prepared for SDS-PAGE and immunoblotting against NRF1 and β-tubulin as the loading control. NRF1 knockdown efficiency (upper band in top panel) was quantified after normalizing with β-tubulin levels using ImageJ, and the normalized values shown. (**C**) Composite plots of FOS-1A7, SMAD2-1B10, Pol2 and chromatin input. Read counts are plotted along a linear x-axis, except between the TSS and TES (transcript end site) of gene bodies (N=7,309 genes), which is shown as a percentage of TSS-TES distance. **(D)** Motif enrichment analysis of ChIP-exo. Cartoons depict models for binding via the cognate motif of the target ssTF or non-cognate binding. Box plots of TPM expression values of target ssTFs associated to antibodies stratified by AUROC value. Results from analysis of 100 putative ssTF binding motifs within each ChIP-exo dataset with >500 peaks (259 datasets in total). We assigned to each ChIP-exo dataset the PWM with the highest AUROC (“Top Motif”) and quantified its centering as the mean distance of the PWM match from the peak’s summit. In the scatter plot, each point represents the enrichment/centering of the top motif in one of the 259 putative TF ChIP-exo datasets. Colors indicate the expression level (RNA-seq TPM value (Consortium 2012); unavailable values are shown in gray) of the gene specific for the antibody used in the ChIP assay. Point sizes indicate the number of ChIP-exo peaks in the dataset. Top motifs with AUROC >0.6 (dashed line) and TPM values from duplicate RNA-seq experiments are indicated. **(E)** Results from enrichment analysis of 100 TF binding motifs within each of 19 ChIP-seq datasets. Points are formatted as in (**D**).

Two mAbs (FOS-1A7 and SMAD2-1B10) gave unexpected ChIP-seq patterns for their intended targets; namely enrichment at transcription start sites (TSS in **Figure 2C**). Previous studies have shown SMAD2 and FOS binding predominately occurs within active enhancers and not at TSS (Aragon et al. 2019; Su et al. 2020). However, both ChIP-seq and ChIP-exo showed enrichment at TSSs and not at predicted (chromHMM) enhancers (see also online data). Deeper biochemical validation will be needed to verify the specificity of these mAbs.

### Assessment through feature enrichment

Thus far we have established the utility of three independent validation criteria inherent to ChIP-exo analysis: 1) enrichment and patterning at a cognate motif (**Figure 1A**); 2) correlation with an independent mAb clone (**Figure 1B**); and 3) co-localization with an interacting partner (**Figure 1C**). In our large-scale evaluation of mAbs, these validation criteria often were either not applicable or not attainable. We therefore looked for additional criteria that might be useful where the preferred validation criteria were inconclusive. We used the ChExMix algorithm to identify significant modes of protein binding and *de novo* motif detection through a combination of DNA sequence enrichment and variation in ChIP-exo patterning (Yamada et al. 2019). Discovered motifs were identified using TOMTOM and the JASPAR database (Gupta et al. 2007; Fornes et al. 2020). Next, their relative enrichment in annotated genomic regions (e.g., promoters, enhancers, and insulators) was quantified. Enrichment may be suggestive of function. These included chromHMM and Segway genome segmentations (*Supplementary Fig. 3A*) (Hoffman et al. 2013).

We also considered a low stringency test that did not require statistical enrichment of peaks (useful for low coverage). Composite plots were generated around well-defined general features like transcription start sites and CTCF binding sites and the average tag enrichment was examined relative to a negative control (IgG) background. We caution that any enrichments for particular targets relative to these features should be followed up with additional replicates. Test results for all validation criteria and other analyses for each tested antibody can be found at www.PCRPvalidation.org and *Supplementary Table 1*. The website provides a deep and rich resource for preliminary discovery for each target, particularly since the vast majority of targets remain uncharacterized (at all, or in the manner(s) we have performed). We caution that the website provides automated analysis for all datasets, including those that did not pass our significance thresholds and/or were not replicated independently. Some of these may simply reflect non-optimized ChIP conditions (e.g., antibody amounts and/or cell type). These additional analyses generally offered less confidence compared to *a priori* cognate motif validation because many of the defined chromatin states are relatively abundant in the genome. The analyses should serve only as a reference point for additional characterization and optimization, and not used to draw biological conclusions.

### Evaluation through motif analysis

To further evaluate the ChIP-exo and ChIP-seq data for evidence of ssTF genomic occupancy (direct or indirect), we analyzed 259 ChIP-exo and 19 ChIP-seq peak files for ssTF motif enrichment by using an Area Under the Receiving Operator Characteristics curve (AUROC) metric in which ChIP ‘bound’ regions are compared to a background set of unbound sequences. Briefly, the AUROC assesses the enrichment of matches to a given TF motif among the ChIP ‘bound’ regions as compared to a background set of unbound regions; the resulting AUROC value ranges from 0 to 1, with 0.5 corresponding to that expected at random. In addition to AUROC motif enrichment, we also quantified the distance of the motif to the peak summit, which is expected to be shorter for motifs recruiting the profiled ssTF to the DNA (either directly or through a tethering ssTF partner) (Gordan et al. 2009; Bailey and Machanick 2012; Wang et al. 2012; Mariani et al. 2017) (**Figure 2C,D**). We used a collection of 100 non-redundant position weight matrices (PWMs) representative of the known repertoire of human ssTF binding specificity (Bailey and Machanick 2012). These approaches identified 20 PCRP antibodies, corresponding to 16 putative ssTFs, for which their cognate DNA motif was both enriched and centered within the ChIP peaks (“Direct Binding” in *Supplementary Table 3*). Possible reasons for the remaining datasets not showing significant motif enrichment include (but are not limited to): (i) the target TF was not expressed at sufficiently high levels or at sufficiently high nuclear concentrations in the assayed cells, (ii) the epitope recognized by the antibody was not accessible in the chromatin context in the assayed cells, (iii) the target TF was not occupying specific genomic target sites (either directly or indirectly) in the assayed cells, or (iv) off-target recognition by the antibody of other proteins in the assayed cells, resulting in lack of sufficient enrichment of the intended target TF.

Of the analyzed datasets, an additional 30 PCRP mAbs showed enrichment for a binding motif other than the cognate motif of the ChIP-profiled ssTF (**Figure 2C,D**). The enrichment of a non-cognate motif suggests that the genomic occupancy of the ChIP-profiled ssTF might be mediated through indirect binding by a different ssTF (Wang et al. 2012; Mariani et al. 2017), which is bound directly to those ChIP ‘bound’ genomic sites through the enriched motif (“Indirect Binding” in *Supplementary Table 3*). In this way, we have previously identified indirect binding modes from ChIP-chip or ChIP-seq experiments that used traditionally prepared antibodies against either an epitope tag on yeast ssTFs or human ssTFs (Gordan et al. 2009; Mariani et al. 2017). Here, for example, the NFAT motifs was enriched and centered among ChIP-seq peaks resulting from ChIP-seq experiments using two different anti-PRDM4 PCRP antibody clones (2B3 and 2B4), suggesting that in HCT116 cells PRDM4 binds DNA indirectly via an NFAT factor. As none of the indirect binding modes that we inferred in this study have been described previously to our knowledge, future experiments are needed to verify them and to rule out the alternative possibility that the antibody may preferentially cross-react with a TF whose motif was found enriched.

### ChIP assessment in multiple cell states and types

Any number of targets may be sequestered in a state that prevents their interaction with chromatin (and thus detection by ChIP) unless activated to do so through a change in cell state. We examined this with HSF1, which is rapidly induced to bind in the nucleus upon heat shock to activate heat shock response genes (Baler et al. 1993). HSF1 was bound to cognate motifs at relatively low levels under non-stressed conditions but increased in binding upon treatment of cells with hydrogen peroxide (0.3 mM) for 3 min and 30 min. (*Supplementary Fig. 5A*, assayed by ChIP-exo), or upon heat shock (shift from 37ºC to 42ºC for 1 hr) (*Supplementary Fig. 5B*, assayed by ChIP-seq). This illustrates a potential problem with using mRNA expression levels as a basis for expecting a factor to be actively bound to chromatin. Many TFs are sequestered and only bind chromatin when released by signaling events.

While ChIP-exo antibody assessments were primarily performed in K562 cells (see above), many targets may have chromatin interactions that are cell-type specific. An example of cell-type specific expression was observed with the breast cancer factor GRHL2, where binding was detected in MCF7 cells but not in other cell types (**Figure 3A**). Other targets like USF1 and NRF1 were less cell type-specific, although we do not exclude selectivity at subsets of sites (**Figure 3B**). Therefore, testing antibodies in the appropriate cell type (with appropriate signaling events) may be critical for target detection.

**Figure 3.**
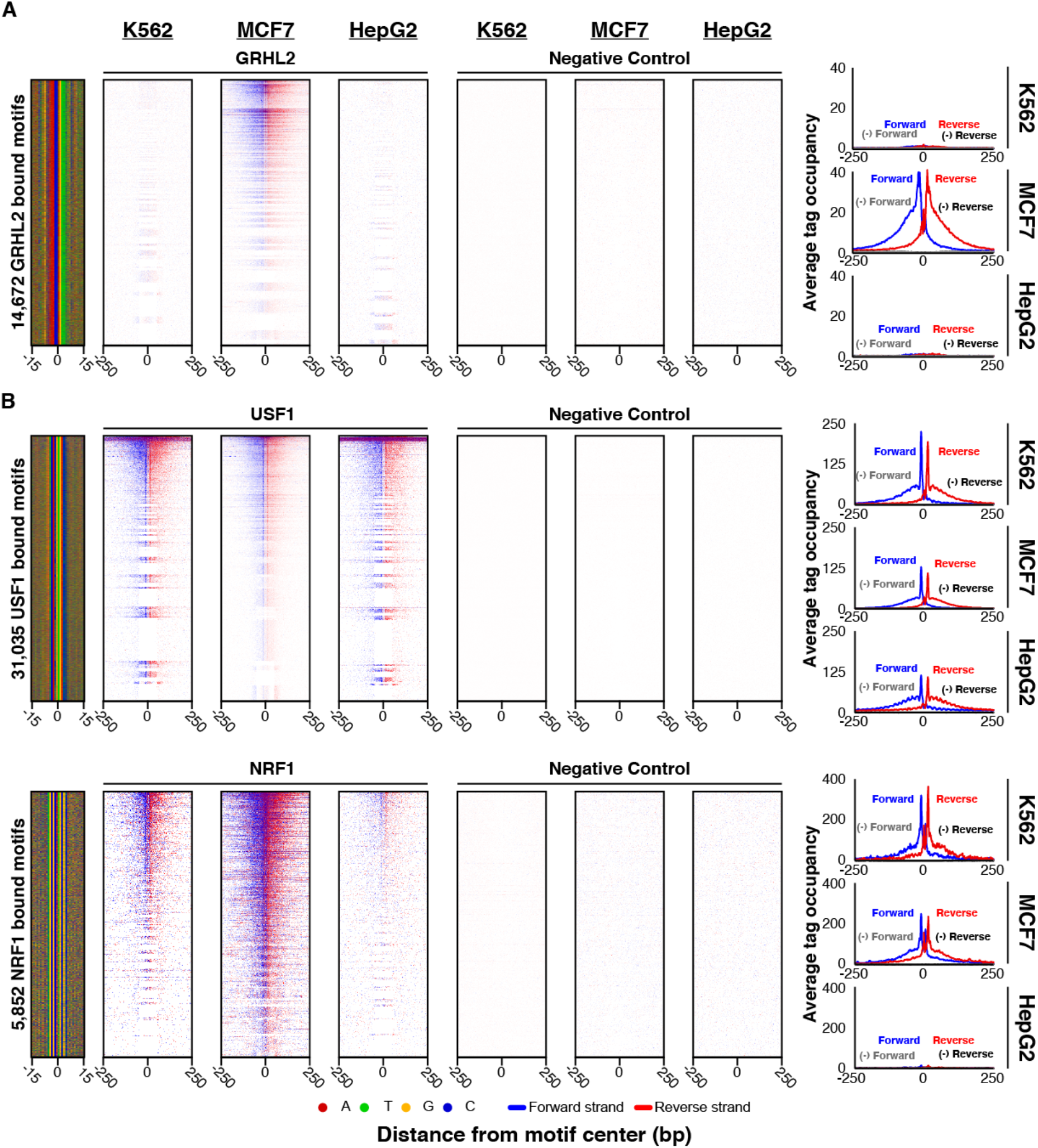
Cell type comparison of antibody performance. (**A**,**B**) ChIP-exo heatmap, composite, and DNA-sequence 4-color plots are shown for the indicated number of bound motifs for the indicated targets, in the indicated cell types. The 5’ end of aligned sequence reads for each set of experiments were plotted against distance from cognate motif, present in the union of all called peaks among the datasets for each indicated target. Reads are strand-separated (blue = motif strand, red = opposite strand). Rows are linked across samples and sorted based on their combined average in a 100 bp bin around each motif midpoint.

We next tested a subset of PCRP mAbs on donated de-identified human organs. ChIP-exo was performed using mAbs against USF1, YY1, and GABPA in chromatin from human liver (2 different specimens), kidney, placenta, and breast tissue (**Figure 4**). Largely consistent with the cell line ChIP’s, enrichment and aligned read patterning was observed with all three mAbs for the liver and kidney, that was diminished in placenta and not detectable in breast. It remains to be determined whether the lack of signal in breast is due to technical limitations in chromatin yields versus tissue specificity of chromatin interactions. Nonetheless, these findings demonstrate the utility of at least some PCRP mAb in epigenomic profiling of human clinical specimens.

**Figure 4.**
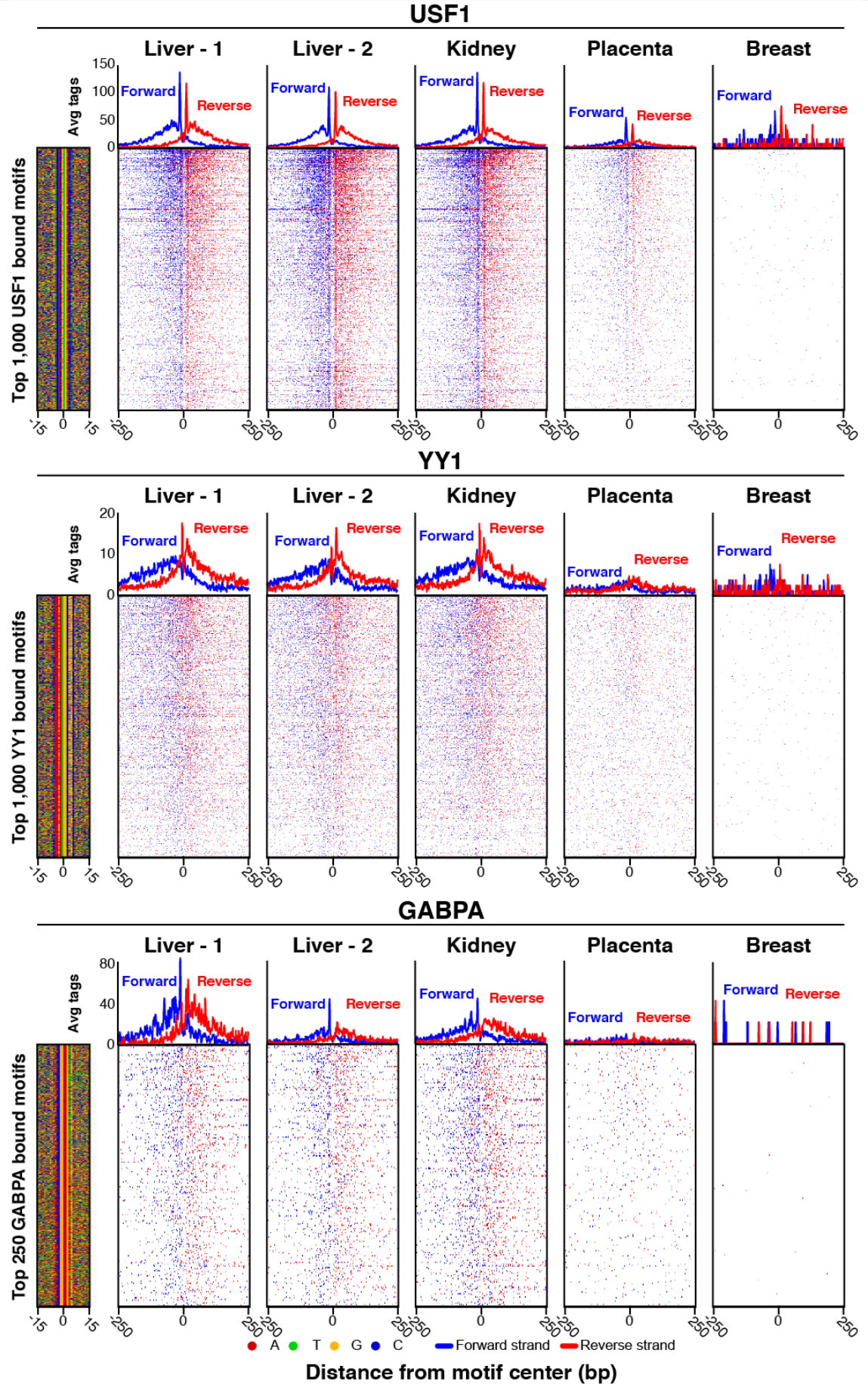
Application of ChIP-exo in human tissue using PCRP mAbs. ChIP-exo heatmap, composite, and DNA-sequence 4-color plots are shown for the indicated number and type of bound motifs for the indicated targets, in the indicated organ types (liver includes two donors). The 5’ end of aligned sequence reads for each set of experiments were plotted against distance from cognate motif, present in the union of all called peaks between the datasets for each indicated target. Reads are strand-separated (blue = motif strand, red = opposite strand). Rows are linked across samples and sorted based on their combined average in a 100 bp bin around each motif midpoint.

### Evaluation using CUT&RUN

CUT&RUN has been used to measure genome-wide protein-DNA interactions (Skene et al. 2018). It uses a fusion of protein A/G (pAG; which binds most antibody isotypes in common use) and micrococcal nuclease (MNase). A ssTF-specific antibody is added to immobilized permeabilized cells or nuclei (under native or crosslinked conditions), where it binds to its chromatin target. pAG-MNase is next added, recruited via pAG to the target-specific antibody, and the MNase portion cleaves local DNA. The result is a selective release of chromatin from the otherwise insoluble nucleus, where genomic enrichment can be identified by sequencing. We tested 40 PCRP antibodies in K562 by native CUT&RUN in replicate (see Methods), of which 25 had been selected based on ChIP-exo enrichment. For USF2, NRF1, USF1, and YY1, we mapped CUT&RUN cleavage sites around their ChIP-exo detected cognate motifs. Multiple nonspecific IgGs served as the negative controls. As additional negative controls, we mapped DNA cleavages around this same set of motifs using the other noncognate ssTF datasets, where only background cleavage is expected (as we did for ChIP-exo in **Figure 1** and ChIP-seq in *Supplementary Fig. 4*). Of all 40 PCRP mAbs antibodies tested, USF2-1A11 produced the most robust CUT&RUN signal (**Figure 5A**), with a detection level matching ChIP-exo, and low IgG-only background. Thus, the native CUT&RUN assay as implemented here has the ability to detect site-specific protein-DNA interactions through at least one PCRP mAb. However, for the NRF1, USF1, and YY1 mAbs, which had worked well in ChIP-exo, we observed little or no enrichment above background in native CUT&RUN (**Figure 5B**). This may reflect intrinsic target incompatibility with the native approach, or that antibody-specific optimization is warranted. Analysis results for the remaining CUT&RUN datasets and controls are in *Supplementary Table 4*. Of note, a DNA-accessibility footprint was observed in some negative control experiments, including a CTCF dataset at noncognate CTCF locations (e.g., at NRF1 and USF1 motifs in **Figure 5B**). This may indicate background cleavage by untargeted pAG-MNase at TF binding sites, perhaps due to the open chromatin at these locations or an intrinsic MNase sequence bias and demonstrates the importance of controls with the CUT&RUN approach. Background cleavage intensity may also vary based on the amount and type of IgG used in the negative control, making peak-calling less reliable. Thus, cognate target specificity in most of these CUT&RUN experiments was not established.

**Figure 5.**
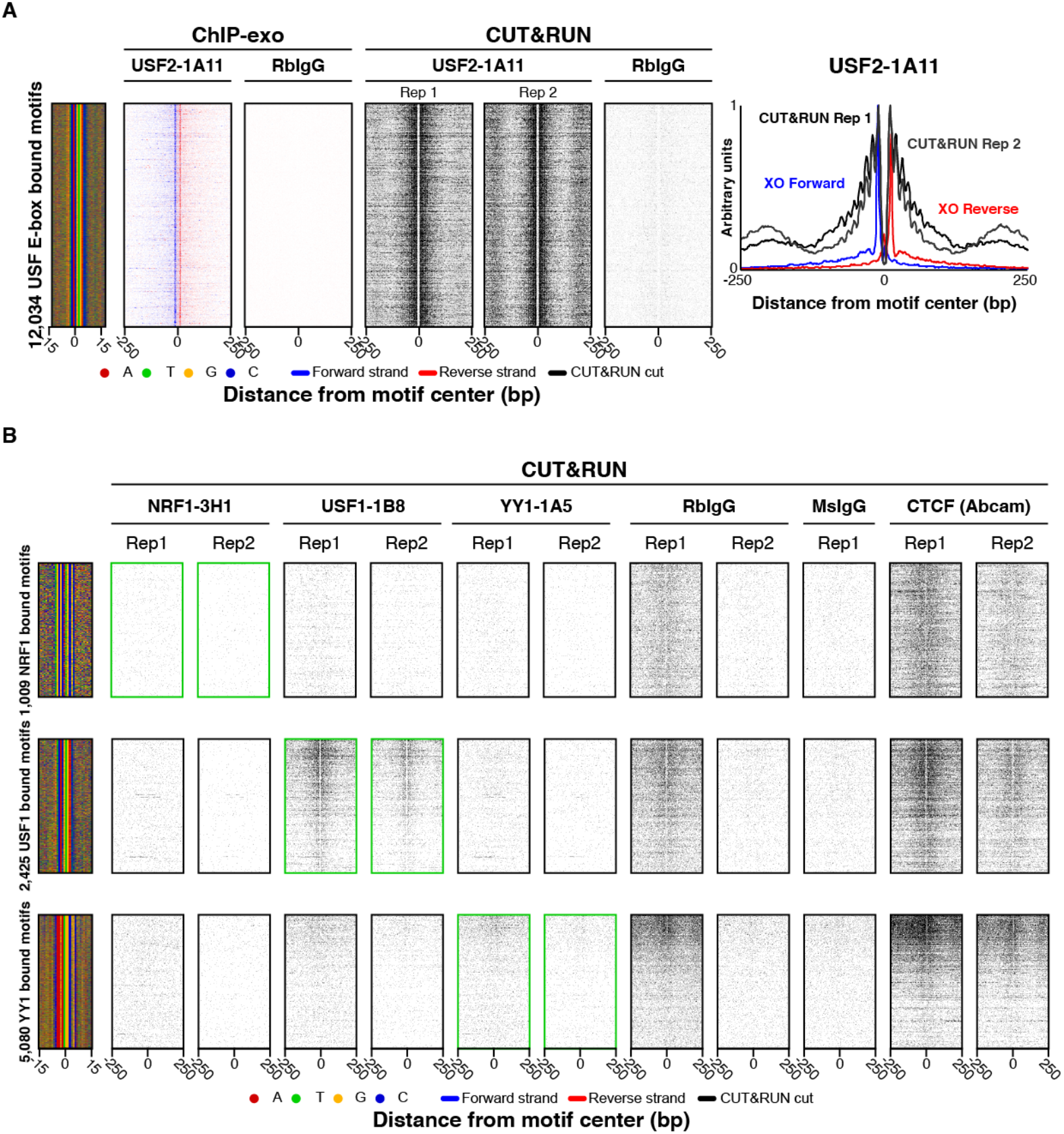
PCRP mAb assayed by CUT&RUN. (**A**,**B**) CUT&RUN heatmap, composite, and DNA-sequence 4-color plots are shown relative to the motifs defined and sorted in Figure 2. The 5’ end of aligned sequence reads are plotted. Reads are strand-separated (blue = motif strand, red = opposite strand) for ChIP-exo and combined (black) for CUT&RUN. Reads are aligned as above (Figure 1A) and individual dataset results are available in *Supplementary Table 4*.

### Evaluation by STORM

As part of our PCRP mAb evaluation, we performed Stochastic Optical Reconstruction Microscopy (STORM), which can visualize cellular structures / processes at nanometer resolution (Betzig et al. 2006). The approach involves the use of fluorescently conjugated antibodies that might be expected to bind identifiable structures in specific subcellular compartments. Of the 39 PCRP mAbs surveyed (*Supplementary Table 5*), most displayed peri-cytoplasmic staining, rather than expected punctate nuclear staining (*Supplementary Fig. 6*). Thus, without further supporting evidence, these results were inconclusive. However, we provide these images as comparison datasets for future studies.

### Evaluation using *in vitro* binding assays

We next tested 44 PCRP mAbs by *in vitro* protein binding assays. A classic method to evaluate antibody specificity is western blotting: size separation of complex protein mixtures using denaturing gel electrophoresis (SDS-PAGE), followed by membrane transfer, and immunoprobing with an antibody of interest to determine the protein species it detects. Since endogenous targets can exist at a level below the sensitivity of detection, we used coupled-in vitro transcription/translation (IVT) in crude HeLa cell extracts to produce 32 TFs as unpurified amino-terminal GST-fusion proteins (*Supplementary Table 6*). This allowed for production of higher levels of target proteins, but within a complex *milieu* of other proteins to allow specificity to be addressed. Of the 44 PCRP mAbs assayed by immunoblotting, 31 (70%) mAbs detected a single predominant band of the expected molecular weight (twelve as biologically independent replicates of the same antibody, plus nine replicated with a different mAb, plus ten performed as a single replicate, *Supplementary Fig. 7*). As a positive control, anti-GST antibody detected 32 of the 33 GST-fusion TFs. Thus, about two-thirds of the assayed PCRP mAbs were specific in recognizing their target proteins. This success rate (70%) may represent the upper limit of success for these reagents.

Protein binding microarrays (PBMs) is a technique to assay protein-DNA binding specificity *in vitro* (Mukherjee et al. 2004; Berger et al. 2006; Siggers et al. 2011b). Proteins used in PBMs are typically expressed as epitope tag-fusions, supporting detection on the DNA array by fluorescent anti-tag antibody. A possible explanation for why some antibodies may fail to work in ChIP experiments is that their target epitope may become inaccessible when the ssTF is bound to a protein partner, DNA ligand, and/or subjected to modification by formaldehyde. Therefore, for a set of 31 ssTFs that were of interest or performed well in ChIP, we used PBMs to test 44 PCRP mAb for their ability to recognize their DNA-bound target TF.

Briefly, the relevant IVT-generated TFs were incubated with DNA microarrays where all possible 10-bp sequences were represented within ∼44,000 60-bp probes on double-stranded oligonucleotide arrays (Agilent) (Berger et al. 2006). For 20 of these 44 PCRP mAbs (45%) assayed against 16 of 22 tested targets, the PBM experiments successfully identified a DNA binding motif consistent with the known or anticipated element (**Figure 6**). All mAb PBM experiments were run with a parallel anti-GST antibody (positive control) to validate the viability of the IVT-expressed target in PBM resulting in a 21/22 (95%) validation rate. Of the 20 PCRP antibodies that successfully yielded the expected motif in PBMs, 11 (55%) had at least some validation support by ChIP.

**Figure 6.**
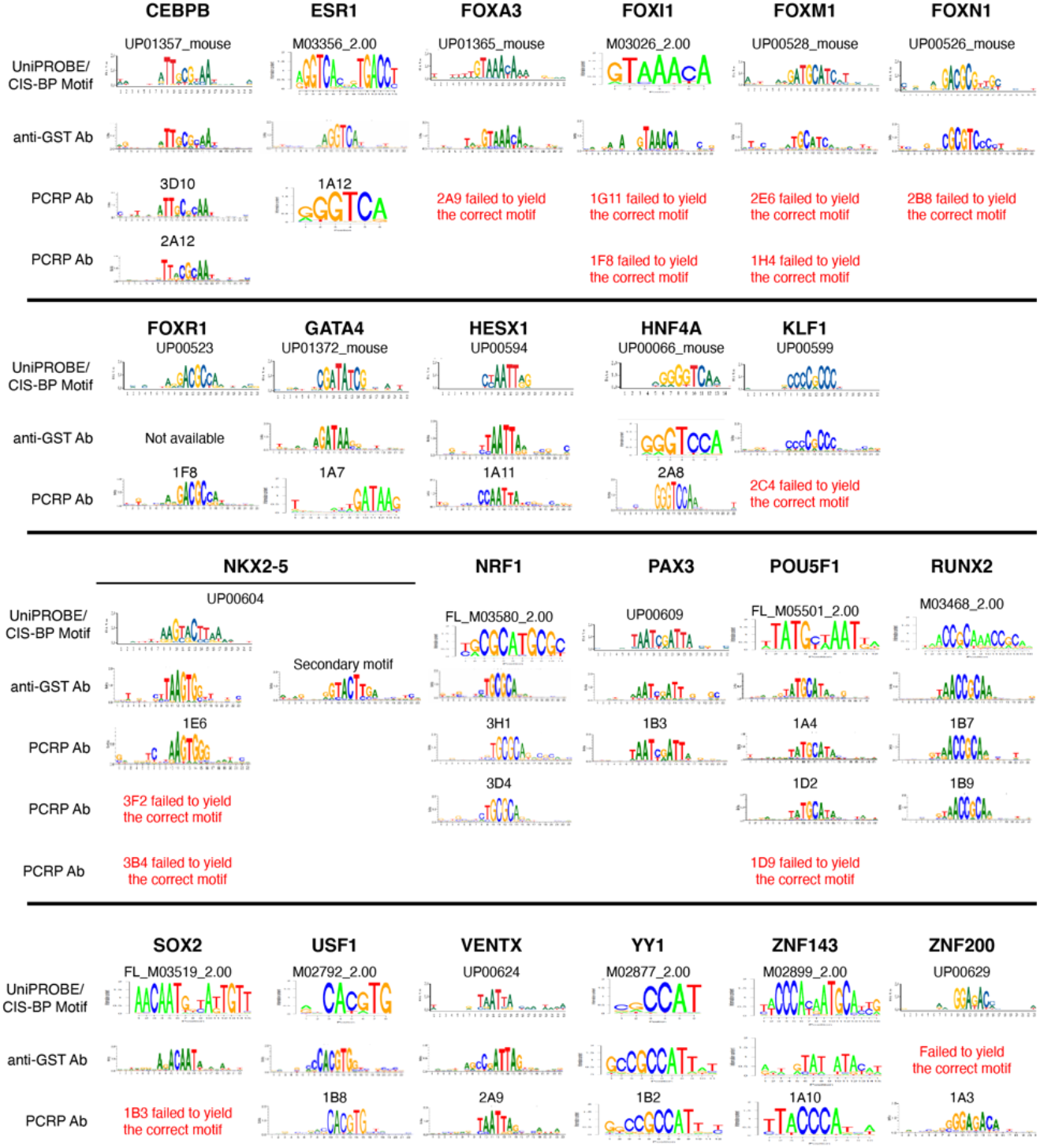
TF binding motifs derived from PBM experiments performed using anti-GST or PCRP antibodies. Each full-length TF was assayed with its PCRP mAb(s) and compared against its corresponding motif derived from an anti-GST PBM experiment and anticipated motif from the UniPROBE or CIS-BP databases (Weirauch et al. 2014; Hume et al. 2015). The TFs from UniPROBE or CIS-BP were assayed as extended DNA binding domains. For display of sequence motifs, probability matrices were trimmed from left and right until 2 consecutive positions with information content of 0.3 or greater were encountered, and logos were generated from the resulting trimmed matrices using enoLOGOS (Workman et al. 2005).

## Discussion

The ability to interrogate the diverse human proteome is heavily reliant on specific affinity capture reagents, of which antibodies are the most widely used. PCRP represented a pilot project for the entire human proteome, with initial focus on nuclear proteins. To this end, this study assayed nearly all PCRP mAb against ∼700 putative chromatin targets or TF using the genome-wide high resolution ChIP-exo assay. A smaller subset was analyzed by other genome-wide assays (ChIP-seq, CUT&RUN); super-resolution microscopy (STORM); to directly detect recombinant target proteins by immunoblotting; or their DNA-binding by PBMs. Our purpose is to present a technical “field” assessment of PCRP mAb utility in biochemical and cellular assays. Given the published rigorous criteria for antibody validation (Landt et al. 2012; Wardle and Tan 2015; Uhlen et al. 2016; Edfors et al. 2018; Sikorski et al. 2018), which may be assay specific, this work is not intended to provide a comprehensive resource of validated antibodies. Instead, it is a starting point for considering validation criteria and any limits that may be applicable to particular assays, especially when taking a systematic high-throughput approach. ChIP-exo identified up to 5% of the ∼1,000 tested PCRP antibodies as having high specificity for their targets, based on orthogonal evidence of motif enrichment and other criteria. These reagents would be the strongest candidates for more rigorous validation testing.

In contrast, a large number of PCRP mAbs did not meet the defined validation criteria we employed. We suggest that some producing ambiguous outcomes might benefit from assay optimization: e.g., a different cell type or growth condition in which the target is expressed and/or activated for chromatin binding. Additionally, different metrics or validity thresholds may be needed. Notably, many of the PCRP mAbs were evaluated by ChIP-exo in K562, while their target TFs may not be appreciably expressed in these cells. However, as shown for HSF1, even where a target is expressed, it may not substantially interact with chromatin (and thus escape detection) unless activated to do so. Therefore, knowledge of the underlying biology of the target may be critical in how ChIP specificity is assessed.

Several algorithmic explanations can be considered for lack of target detection. Some sequence-specific DNA binding proteins may not have been accommodated within our discovery framework. For example, a target protein may bind a nonstandard distribution DNA sequences that was not captured by current motif discovery algorithms. Alternatively, the target may interact with a wide range of genomic sites having different or degenerate DNA sequence motifs (including indirect sequence readout based on DNA shape (Rohs et al. 2009)) that are not accommodated by the discovery algorithms used in this study. Another possible scenario is that a target protein might not bind DNA directly, but only indirectly through other proteins, including those that potentially form an undiscovered chromatin class (which would not be in our discovery pipeline). We often found motifs that were long, simple, semi-repetitive, and highly degenerate (see online). These are not typical properties of sequence-specific DNA binding proteins and the ChIP-exo patterns at these motifs were often quite distinct from well-validated targets. Historically, some of these locations may have been set aside as problematic, and thus excluded from analysis (Consortium 2012). Whether these regions are artifactual or have some unknown biology remains to be determined. While we accepted these motifs as evidence of enrichment, we urge caution when interpreting such atypical binding events.

It was not practical in our high-throughput ChIP-exo screen to profile each PCRP mAb in a wide range of cell types. However, for ssTFs with at least 500 significant peaks, we noticed an association between the expression of the target proteins and the detection of binding motif specificity (box plot in **Figure 2D**), and so expression may be a useful preliminary guide for cell-type selection. Furthermore, some targets may simply not be cross-linkable to chromatin in the assayed cell type (or any cell type), making ChIP an inappropriate assay. Unlike engineered epitope tags, each target-specific antibody may have a substantially different affinity for its cognate antigen. Therefore, we cannot rule out that at least some low- or non-performing antibodies could perform better under different immunoprecipitation conditions. Still other potential reasons for antibody non-performance may be due to trivial explanations like lot expiration or mis-labeling along the supply chain.

In total, 946 unique hybridoma clones were tested in at least one of the assays. We identified 50 clones (5%) that worked with high confidence in at least one assay. However, only a very small portion of the validation spectrum has been explored. Using relaxed criteria, that may reflect significant but off-target or unknown behavior, we find that 371 (39%) of the tested PCRP mAb had at least some evidence of being different from background, in at least one assay. However, such marginal criteria require deeper characterization, such as target depletion/deletion or negative control cell lines for a more robust validation. The remaining 61% also warrant more testing in other cell types and conditions. Our analysis identifies an initial set of prioritized candidates. A detailed summary of each assay’s results along with all of the measured quality control metrics is available in **Table 1**, along with an interactive searchable web-interface online at www.PCRPvalidation.org.

## Methods

### ChIP protocols

#### Antibodies

1,308 TF hybridomas were reported through the PCRP portal at the start of this study (September 2017). Hybridoma supernatants were purchased from Developmental Studies Hybridoma Bank (DSHB, U. Iowa, IA) as 1 ml aliquots. Monoclonal antibody (mAb) concentration averaged 36 ug/ml by ELISA quantification. Hybridoma supernatants contain ADCF-MAb cell culture medium (https://dshb.biology.uiowa.edu/tech-info.) and residual (2%) fetal bovine serum, which has a reported IgG concentration of 1-6 ug/ml (Son et al. 2001). DSHB preparation dates were provided. Concentrated mAbs and their concentration were generously provided by CDI laboratories (Mayaguez, PR).

#### Cell material

Cell stocks were obtained by the Pugh laboratory from ATCC. K562 were grown in suspension using IMDM media and periodically checked for mycoplasma contamination. HepG2 and MCF7 were grown as adherent cells in DMEM media. MCF7 cells were additionally grown in phenol red-free DMEM and treated with Beta-estradiol 30 min prior to cell harvest. Cells were pelleted, re-suspended in PBS, crosslinked with 1% formaldehyde for 10 min, and then quenched with a molar excess of glycine. Donated human organs were obtained from NDRI (Philadelphia, PA) and then cryoground to fine powder using the SPEX cryomill cyrogrinder. Frozen tissue powder was re-suspended in room-temperature PBS containing formaldehyde to a final concentration of 1% and quenched with a molar excess of glycine. All cells and tissue for ChIP-exo then proceeded through standard lysis and sonication protocol described below after crosslink quenching. HCT116 cells (ATCC CCL-247) were grown in the Shilatifard laboratory in DMEM supplemented with 10% FBS (Fisher Scientific, 35-015-CV). 70%–80% confluent HCT116 cells were heat shocked for 1 hr by adding pre-heated conditioned media pre-heated to 42°C (Lim et al. 2017). Heat shock and non-heat shock HCT116 cells were washed with PBS before fixing with 1% formaldehyde (Sigma, 252549) in PBS for 15 minutes and processing for ChIP-seq.

ChIP-exo testing was initially prioritized in K562, MCF7, and HepG2, using gene expression values (FPKM in RNA-seq) generated from the ENCODE project as the basis for the cell type used (Uhlen et al. 2005; Consortium 2012). Targets were assigned to the cell type most likely to express the protein of interest. If no cell line had a clear high expression for a specific target (>25% FPKM relative to all other considered cell lines), testing defaulted to K562. K562 was selected as the default due to its status as a Tier 1 ENCODE cell line and the plethora of existing genomic data that could orthogonally support any findings. Samples were processed in batches of 48 in 96 well plate format. The PCRP-derived USF1 or NRF1 antibody served as a positive control for every processed cohort as well as an IgG or “No Antibody” mock ChIP negative control. Crosslinked sonicated chromatin from ∼7 million cells was incubated with antibody-bound beads, then subjected to the ChIP-exo 5.0 assay (Rossi et al. 2018).

#### ChIP-exo 5.0 assay

Chromatin for ChIP-exo was prepared by resuspending crosslinked and quenched chromatin in Farnham cell lysis buffer at a ratio of 25 million cells to 1 mL of buffer for 20 min at 4°C. At the 10 min mark, cells were pushed through a 25G needle 5 times to enhance cellular lysis. Nuclei were then isolated by pelleting at 2,500g for 5min. Nuclei were resuspended in RIPA buffer (25 million cells to 1 mL of buffer) for an additional 20 min at 4°C and then pelleted again at 2,500g for 5 min. Disrupted nuclei were then finally resuspended in 1X PBS (25 million cells to 1 mL of buffer) and sonicated for 10 cycles (30on/30off) in a Diagenode Pico. Solubilized chromatin was then processed through ChIP-exo. Production-scale ChIP-exo 5.0 was generally performed in batches of 48 in a 96-well plate, alternating every column to reduce risk of cross-contamination. Briefly, solubilized chromatin was incubated with Protein A/G Dynabeads, preloaded with 3 ug of antibody, overnight then sequentially processed through A-tailing, 1^st^ adapter ligation, Phi29 Fill-in, Lambda exonuclease digestion, cross-link reversal, 2^nd^ adapter ligation, and PCR for final high-throughput sequencing. Equal proportions of ChIP samples were barcoded, pooled, and sequenced. Illumina paired-end read (40 bp Read 1 and 36 bp Read 2) sequencing was performed on a NextSeq 500 and 550. While, on average, we sought ∼10 million total paired-end reads per ChIP, we accepted less if there was strong evidence of target enrichment. Otherwise, we performed an additional round of sequencing. The 5’ end of Read_1 corresponded to the exonuclease stop site, located ∼6 bp upstream of a protein-DNA crosslink. Read_2 served two indirect functions: to provide added specificity to genome-wide mapping, and to remove PCR duplicates.

#### ChIP-seq

1×10^8 cells were fixed in 1% formaldehyde (Sigma, 252549) in PBS for 15-20 min at room temperature and quenched with 1/10th volume of 1.25 M glycine for 5 minutes at room temperature. Cells were collected at 1,000xg for 5 minutes, washed in PBS, pelleted at 1,000xg for 5 min and pellets were flash frozen in liquid nitrogen and stored at −80°C until use. Pellets were thawed on ice and resuspended in 10 ml lysis buffer 1 (50 mM HEPES, pH 7.5, 140 mM NaCl, 1 mM EDTA, 10% glycerol, 0.5% IGEPAL CA-630, 0.25% Triton X-100 with 5µl/ml Sigma 8340 protease inhibitor cocktail incubated on ice 10 minutes, pelleted 1,500xg and subsequently washed in lysis buffer 2 (10 mM Tris-HCl pH 8.0, 200 mM NaCl, 1 mM EDTA, 0.5 mM EGTA and 5ul /ml protease inhibitor) as with lysis buffer 1 before resuspending in 1 ml lysis buffer 3 (10 mM tris-HCl pH 8.0, 1 mM EDTA, 0.1% SDS and 5 ul/ml protease inhibitors) for sonication as previously described (Lee et al. 2006). Chromatin was sheared in a 1 ml milliTUBE with AFA fiber on a Covaris E220 using 10% duty facf”blator for 2 min. The sheared chromatin concentration was estimated with Nanodrop at OD 260 and diluted to 1 mg/ml in ChIP Dilution Buffer (10% Triton X-100, 1M NaCl and 1% Sodium deoxycholate). 1 mg chromatin was combined with 4 µg hybridoma tissue culture supernatant and rotated overnight at 4°C. 40 µl of protein G Dynabeads was added and incubated for 2-4 hr rotating at 4°C. Samples were washed 5 times with 1 ml RIPA buffer, 2 times with TE with 50 mM NaCl. Chromatin was eluted with 800 µl elution buffer (50 mM Tris pH 8.0, 1 mM EDTA, 0.1% SDS) for 30 min at 65°C shaking at 1,500 rpm in a ThermoMixer (Eppendorf). Supernatants were collected, digested with 20 µl of 20 mg/ml Proteinase K and incubated overnight at 65°C. DNA was purified with phenol chloroform extraction. 500 µl of the aqueous phase was precipitated with 20 µl 5M NaCl, 1.5 µg glycogen and 1 ml EtOH on ice for 1 hr or at −20°C overnight.

Sequencing libraries were prepared with the KAPA HTP library prep kit (Roche) using 1-10 ng DNA and libraries were size selected with AMPure XP beads (Beckman Coulter). Illumina 50 bp single-end read sequencing was performed on a NextSeq 500 or NovaSeq 6000. The modular pipeline Ceto (https://github.com/ebartom/NGSbartom) was used to convert base calls to fastq, align reads to bam files and make bigWig coverage tracks. Briefly, bcl2fastq with parameters -r 10 -d 10 -p 10 -w 10 was used to generate fastq files. Trimmomatic version 0.33 with the options single end mode (SE) and -phred33 was used to remove low-quality reads. Reads were then aligned to hg19 with Bowtie 1.1.2 (Langmead et al. 2009) with options -p 10 -m 1 -v 2 -S, thus keeping only uniquely mapped reads, and allowing up to two mismatches. Coverage tracks were created with the Rscript from Ceto, createChIPtracks.R --extLen=150 to extend reads to 150 bp, and coverage was normalized to total mapped reads (reads per million). Peaks were called with MACS2 2.1.0 (Zhang et al. 2008) with a cutoff of -q 0.01 and the input chromatin used as the control dataset. Heatmaps and composite plots were made with deepTools (Ramirez et al. 2016) version 2.0. computeMatrix using reference-point peaks. “Black-listed” regions were removed with parameter -bl Anshul_Hg19UltraHighSignalArtifactRegions.bed (ftp://encodeftp.cse.ucsc.edu/users/akundaje/rawdata/blacklists/hg19). For metagene plots, we used the Ensembl version 75 transcripts with the highest total coverage from the annotated TSS to 200 nt downstream and were at least 2 kb long, and 1 kb away from the nearest gene.

#### RNAi

Lentiviruses were packaged in HEK293T cells transfected with 1 µg pME-VSVG, 2 µg PAX2 and 4 µg shRNA in pLKO.1 backbone using Lipofectamine 3000 (Thermo Fisher Scientific, Waltham, MA) according to the manufacturer’s instructions. The virus particles were then harvested 24 to 48 hours later by passing through a 0.45 µm syringe filter (Thermo Scientific). Viruses were mixed with an equal volume of fresh media supplemented with 10% FBS and polybrene was added at a final concentration of 5 µg/mL to increase infection efficiency. The medium was changed 6 hours after infection. Cells were selected with 2 µg/ml puromycin for three days before Western blotting (anti-NRF1 rabbit mAb clone D9K6R from CST and anti-beta tubulin mouse mAb clone E7 from DSHB) and ChIP-seq experiments. The following shRNA sequences were used:

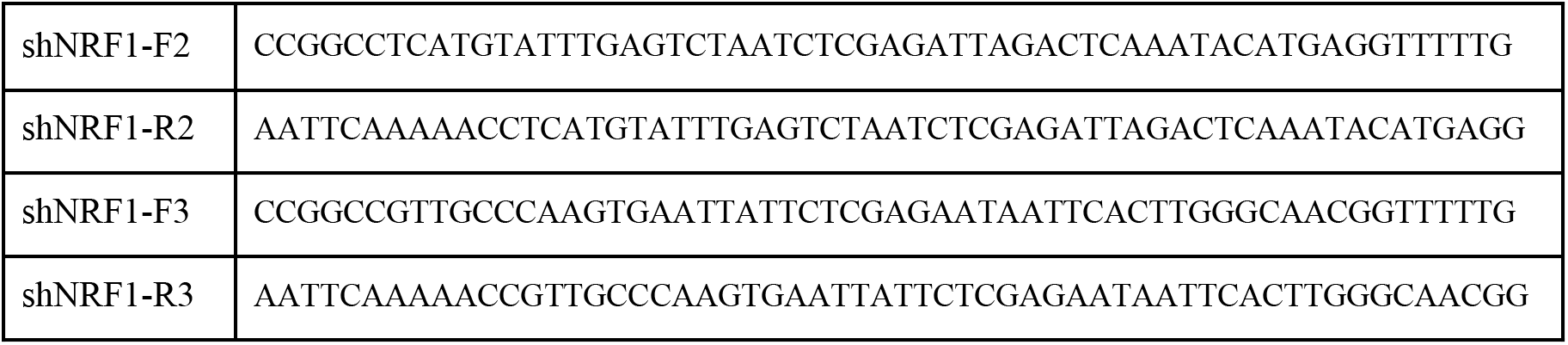

### Bioinformatic protocols

#### Technical performance

A series of modular bioinformatic analyses were implemented to evaluate technical success of ChIP-exo library construction and sequencing, independent of whether the antibody found its target or not. 1) Sequencing depth (standard is 8-10M), which is the total number of sequencing reads having a target-specific barcode. 2) Adapter dimers (standard is <2%), which is the fraction of reads that contain the sequencing adapters but lack a genomic insert. 3) Alignment (standard is 70-90%), which is the percent of reads that map to the reference human genome after removing adapter dimers. 4) PCR duplicates (standard is <40%), which are expected to have identically mapped Read_1 and Read_2 5’ ends. We assume that when Read_1 and Read_2 5’ ends have identical mapped coordinates; they represent PCR duplicates. Since Read_2 is generated by sonication it is expected to be distributed across a region, and thus not likely to be at the same coordinate twice. Since PCR duplicates are not a direct product of ChIP, they add no value to enrichment metrics. High PCR duplicates, at normal sequencing depths, often means technical loss of material during library construction prior to PCR.

#### Peak calling

We utilized two distinct algorithms for ChIP-exo peak-calling. The first algorithm was GeneTrack which used a gaussian kernel to call strand-separated peaks at the 5’ ends of reads (Albert et al. 2008). The reads were then paired across strands and the tag occupancy was normalized using the NCIS approach (Liang and Keles 2012). Peak significance was called using either the binomial or Poisson test (taking whichever p-value was higher) with Benjamini-Hochberg correction and a q-value cutoff of q<0.01. GeneTrack was used to generate the ChIP-exo peaks used in the manuscript figures. The other peak-caller used was the ChExMix algorithm, which is a high-resolution peak-caller designed to simultaneously identify enriched sequence motifs and distinct sub-types of binding using a combination of clustering and hierarchical mixture modeling (Yamada et al. 2019). ChExMix was designed to take advantage of ChIP-exo’s ability to identify protein co-factors through indirect crosslinking events by modeling detected tag distributions in ChIP-exo data and using tree-clustering based approaches to determine the significant peak subtypes that exist with ChIP-exo data. ChExMix-called peaks were used to interrogate the ChIP-exo peaks subtype structure and are visualized on the www.PCRPvalidation.org website.

#### Motif enrichment via ChExMix

De novo motif discovery was performed by ChExMix. Each motif was compared against the JASPAR database using TOMTOM with default parameters to identify similarity to known motifs (Bailey et al. 2009; Fornes et al. 2020). Heatmaps and composite plots were generated of sequence reads aligned relative to motif midpoints of all peaks containing an enriched motif. For samples with high background, low complexity, and/or low sequencing depth, it is possible the antibody is valid, but that standard de novo motif discovery may fail. We developed an orthogonal method for motif detection. By first identifying all non-redundant motif classes (Castro-Mondragon et al. 2017) in the genome, we then overlap low-threshold ChExMix peaks and determined which motif class possesses overlapping peaks above background (>2 log_2_).

#### Motif enrichment and centering analysis

For ChIP-exo data generated from K562, HepG2 and MCF-7 cells, narrowpeak data were called using ChExMix (Yamada et al. 2019); we restricted our motif enrichment analysis to narrowpeak datasets that contained more than 500 peaks. For ChIP-seq data generated from HCT116 cells, we required the presence of 100 peaks due to the typical lower number of called peaks in those datasets as compared to ChIP-exo. Motif enrichment analysis of ChIP-exo and ChIP-seq peaks was then performed as described previously (Mariani et al. 2017). For ChIP-exo peaks, we first filtered for the datasets that had more than 500 peaks, and then used for the comparison the top 500 peaks, with peaks defined as the ChIP-exo summits computationally padded with the region spanning [-100 bp, +100 bp]. To perform an analogous analysis on ChIP-seq peaks, we fixed both the number of peaks per dataset (e.g., top 100 peaks) and the peak size, which we computationally trimmed similarly to the ChIP-exo data to span [-100 bp, +100 bp] surrounding the ChIP-seq peak summit. For each ChIP peak set, we generated background sequences using GENRE software with the default human setting, to ensure the same level of promoter overlap, repeat overlap, GC content and CpG dinucleotide frequency between each peak and its associated background sequence (Mariani et al. 2017). We manually curated a collection of 100 position weight matrices (PWMs), primarily from biochemical TF DNA binding assays (*i.e.*, PBM or HT-SELEX), from the UniPROBE and CisBP databases, as a representative repertoire of human sequence-specific TF binding motifs (Weirauch et al. 2014; Hume et al. 2015; Mariani 2020). We scored each sequence for matches to each of the motifs using the function “matchPWM” from the “Biostrings” R package. Motif enrichment was quantified using an established AUROC metric that assesses the presence of a motif among the 500 highest confidence peaks (foreground set) as compared to the corresponding background set of sequences using publicly available tools for analysis of TF ChIP-seq data (Mariani et al. 2017). We also assessed each motif for its enrichment towards the centers of each ChIP-exo or ChIP-Seq peak set as described previously (Mariani et al. 2017). Briefly, we first identified the PWM score threshold that maximized the difference between foreground and corresponding background sets in the number of sequences containing at least one PWM match (*Optimal PWM Match Score*). If a sequence had multiple PWM matches, we considered only the highest score site. We then calculated the distance from each of these sites to the corresponding peak summits and used the mean of these distances in the foreground or background set to quantify the motif enrichment towards the centers of ChIP peaks. The *P*-values associated with motif enrichment (*i.e.*, AUROC value) and enrichment towards the peak summits (*i.e.*, mean motif distance to peak summit) were both calculated by using a Wilcoxon signed-rank test comparing their scores (PWM match score and PWM match distance to distance to peak summit, respectively) for foreground and background sequences when the PWM threshold was set to the optimal PWM match score. We then adjusted the *P*-values across the PWM collection with a false discovery rate test for multiple hypothesis testing. To test the significance of the difference in the TPM distributions between ChIP datasets with enriched versus non-enriched motifs, we calculated the *P*-value by a Wilcoxon test using the function wilcox.test in R.

#### Genome annotation enrichment

Only a small fraction of DNA-interacting factors binds sequence-specific motifs. In the case of targets with either no expected motif or no known function, determining peak enrichments at annotated regions of the genome can provide evidence of ChIP success. The relative frequency of peaks occurring in different functional genomic regions as defined by chromHMM was calculated for each target, an IgG negative control, and for random expectation (Ernst et al. 2011). The log_2_ frequency enrichment of sample over IgG control was used to identify regions of enrichment, as well as significant areas of de-enrichment (regions that selectively avoid the target). Significant peaks were intersected with chromHMM and Segway states to generate frequency histograms for overlap with predicted chromatin states (Ernst et al. 2011; Hoffman et al. 2012). Peaks derived from the matched negative control dataset were also intersected with annotated states. The log_2_ ratio of sample state frequency over control state frequency was then calculated in order to identify general state enrichment of the sample throughout the genome.

#### Positional enrichment at promoters and insulators

In order to identify enrichment in well-characterized promoter regions, sequence reads for the target, a matched “No Antibody” control, and an IgG were aligned relative to annotated transcription start sites (TSS). Heatmaps of all genes and composite plots of the top 1,000 TSS by gene expression (RNA-seq FPKM) were generated from the data^7^.

#### Heatmaps, composite plots, and 4color sequence plots

All heatmaps, composite plots, and 4 color sequence plots were generated using ScriptManager v0.12 (https://github.com/CEGRcode/scriptmanager). ScriptManager is a Java-based GUI tool that contains a series of interactive wizards that guide the user through transforming aligned BAM files into publication-ready figures.

### CUT&RUN protocols

#### Antibody sourcing and concentration

Antibody hybridoma supernatants, name, clone ID, and lot) were from DSHB. mAbs were concentrated using Amicon Ultra-4 Centrifugal Filter Units with a 50 kDa cut-off (Millipore Sigma Cat # UFC805024) following manufacturer’s recommendations. All centrifugation steps (including 3x 4ml washes with 1X Tris buffered Saline [TBS]) were performed at 4,000 x g for 15 minutes at room temperature. Final concentrations for recovered mAbs (stored at 4°C in [TBS, 0.1% BSA, 0.09% Sodium Azide]) were assumed based on initial concentrations / final recovery volumes and 1 µg used per CUT&RUN experiment.

#### CUT&RUN

CUT&RUN was performed on 500k native nuclei extracted from K562 cells using CUTANA^®^ protocol v1.5.1 [http://www.epicypher.com] which is an optimized version of that previously described (Skene et al. 2018). For each sample, nuclei were extracted by incubating cells on ice for 10 min in Nuclei Extraction buffer (NE: 20 mM HEPES–KOH, pH 7.9; 10 mM KCl; 0.1% Triton X-100; 20% Glycerol; 0.5mM spermidine; 1x complete protease inhibitor [*Roche* # 11836170001]), collecting by centrifugation (600 g, 3 min, 4°C), discarding the supernatant, and resuspending at [100 µl / 500K nuclei] sample in NE buffer. For each target 500K nuclei were immobilized onto Concanavalin-A beads (*EpiCypher* #21-1401) and incubated overnight (4°C with gentle rocking) with 1 µg of antibody (For all 40 PCRP antibodies as above; RbIgG (*EpiCypher* 13-0042, lot 20036001-52); MsIgG (Invitrogen 10400C, lot VD293456); CTCF (*Millipore* 07-729, lot 3205452).

#### Modified CUT&RUN library prep

Illumina sequencing libraries were prepared from 1ng to 10ng of purified CUT&RUN DNA using NEBNext Ultra II DNA Library Prep Kit (*New England Biolabs* # E7645) as previously (Liu et al. 2018) with the following modifications to preserve and enrich smaller DNA fragments (20-70 bp). Briefly, during end repair the cycling time was decreased to 30 mins at 50°C. After adapter ligation, to purify fragments >50bp, 1.75x volumes of Agencourt AMPure XP beads (*Beckman Coulter* #A63881) were added for the first bead clean-up before amplification following manufacturer’s recommendations. PCR amplification cycling parameters were as described (Skene et al. 2018). Post-PCR, two rounds of DNA size selection were performed. For the first selection, 0.8x volume of AMPure XP beads was added to the PCR reaction to remove products >350bp. The supernatant, containing fragments <350bp, was moved forward to a second round of size selection using 1.2x volumes of AMPure XP beads, to remove products <150 bp. Libraries were quantified using Qubit Fluorometer (*Invitrogen*) and checked for size distribution with a Bioanalyzer (*Agilent*).

#### CUT&RUN library sequencing and data analysis

Libraries were sequenced on the *Illumina* NextSeq 550, obtaining ∼5 million paired-end reads (75 x 75 nucleotides) on average. Paired-end FASTQ files were aligned to the hg19 reference genome using the ChIP-exo pipeline.

### TF cloning, protein expression, western blots, and PBM protocols

Full-length TFs were either obtained from the hORFeome clone collection or synthesized as gBlocks (Integrated DNA Technologies) (**Supplementary Table 7**), full-length sequence-verified, and transferred by Gateway recombinational cloning into either the pDEST15 (ThermoFisher Scientific) or pT7CFE1-NHIS-GST (ThermoFisher) vectors for expression as N-terminal GST fusion proteins (Collaboration 2016). TFs were expressed by a coupled *in vitro* transcription and translation kit according to the manufacturer’s protocols (**Supplementary Table 6**). Protein concentrations were approximated by an anti-GST western blot as described previously (Berger et al. 2006). All PCRP antibodies were used at a final concentration of 40 ng/mL in western blots; based on successful outcomes in PBM experiments, PCRP antibodies 1A7 (anti-GATA4), 2A4 (anti-HNF4A), and 1B3 (anti-PAX3) were also used at a final concentration of 1,000 ng/mL in western blots. 8×60K GSE ‘all 10-mer universal’ oligonucleotide arrays (AMADID #030236; Agilent Technologies, Inc.) were double-stranded and used in PBM experiments essentially as described previously, with minor modifications as described below (Berger et al. 2006; Berger and Bulyk 2009; Nakagawa et al. 2013). GST-tagged TFs assayed in PBMs were detected either with Alexa488-conjugated anti-GST antibody (Invitrogen A-11131), or with a TF-specific PCRP antibody, followed by washes and detection with Alexa488-conjugated goat anti-mouse IgG(H+L) Cross-Adsorbed Secondary Antibody (Invitrogen A-11001), essentially as described previously (Siggers et al. 2011b) (**Supplementary Table 6**). All PCRP Abs were used undiluted in PBM experiments; a subset of the PCRP Abs were also tested at a 1:5 or 1:20 dilution (**Supplementary Table 6**). All PBM experiments using PCRP antibodies were performed using fresh arrays or arrays that had been stripped once, as described previously (Berger et al. 2006; Berger and Bulyk 2009). PBMs were scanned in a GenePix 4400A Microarray Scanner and raw data files were quantified and processed using the Universal PBM Analysis Suite (Berger et al. 2006; Berger and Bulyk 2009).

### STORM protocols

#### Supernatant concentration

3 milliliters of PCRP supernatant were concentrated using the Amicon® Pro Affinity Concentration Kit Protein G with 10kDa Amicon® Ultra-0.5 Device following the manufacturer’s recommendations. Supernatant was quantitated by spectrophotometer for rough approximation of concentration.

#### Cellular preparation and staining

K562 cells obtained from the ATCC (cat number CCL243) were grown in a humidified 5% CO2 incubator. 3-5×10^5^ cells were centrifuged at 1,500 rpm for 5 minutes, washed with PBS, and plated on MatTek-brand glass bottom dishes (P35-G.1) prepared appropriately (washes with increasing concentrations of ethanol, followed by coating with poly-L-Lysine (Sigma P4707) for 5 minutes and subsequent washes with water and airdrying for 2h.) After plating, the cells were allowed to adhere for 2h and then washed with PBS. For fixation, 1ml of 4% paraformaldehyde and 0.1% glutaraldehyde in PBS were added for 10 minutes at room temperature with gentle rocking followed by blocking and permeabilization in 2% normal goat serum/1% Triton X-100 in PBS for 1h at room temperature with gentle shaking. Immunostaining was performed with anti-MTR4 antibody (Abcam 70551) at a dilution of 1:250 in 0.1% normal goat serum, 0.05% Triton X-100 overnight at 4 degrees C. The next morning cells were incubated with the secondary antibody (conjugated to Alexa-Fluor 647 (Life Technologies)) at a 1:1000 dilution in 0.1% normal goat serum in PBS and incubated for 2h at room temperature followed by washes in PBS. This material was then subjected to immunostaining with PCRP antibodies, by repeating the above procedure. PCRP mAb hybridoma supernatant (3 ml) was first concentrated 30-100 fold since raw supernatants were unsuccessful in both confocal microscopy and STORM (data not shown). The secondary antibody was conjugated to Alexa-Fluor 488 (Life Technologies). At the end of the application of the second secondary antibody, the cells were washed 3x with PBS and DAPI (Sigma D8542) at a concentration of 1:500 in 0.1% normal goat serum in 1x PBS was applied for 10 minutes at room temperature. The cells were washed 3x with PBS and finally 4% paraformaldehyde/0.1% glutaraldehyde in PBS was applied for 10 minutes at RT before 3 washes in 1x PBS were performed and dishes stored at 4 degrees until microscopy could be performed.

#### Microscopy

Cells were brought to the imaging facility and OXEA buffer was applied (50 mM Cysteimine, 3% v/v Oxyfluor, 20% v/v sodium DL Lactate, with pH adjusted to approximately 8.5, as necessary.) The two colors (Alexa fluor 488 and Alexa fluor 647) were imaged sequentially. Imaging buffer helped to keep dye molecules in a transient dark state. Subsequently, individual dye molecules were excited stochastically with high laser power at their excitation wavelength (488 nm for Alexa fluor 488 or 647 nm for Alexa fluor 647, respectively) to induce blinking on millisecond timescales. STORM images and the correlated high-power confocal stacks were acquired via a CFI Apo TIRF 100 × objective (1.49 NA) on a Nikon Ti-E inverted microscope equipped with a Nikon N-STORM system, an Agilent laser launch system, an Andor iXon Ultra 897 EMCCD (with a cylindrical lens for astigmatic 3D-STORM imaging) camera, and an NSTORM Quad cube (Chroma). This setup was controlled by Nikon NIS-Element AR software with N-STORM module. To obtain images, the field of view was selected based on the live EMCCD image under 488-nm illumination. 3D STORM datasets of 50,000 frames were collected. Lateral drift between frames was corrected by tracking 488, 561, and 647 fluorescent beads (TetraSpeck, Life Technologies). STORM images were processed to acquire coordinates of localization points using the N-STORM module in NIS-Elements AR software. Identical settings were used for every image. Each localization is depicted in the STORM image as a Gaussian peak, the width of which is determined by the number of photons detected (Betzig et al. 2006). All of the 3D STORM imaging was performed on a minimum of two different K562 cells.

### Code Availability

Our automated bioinformatic ChIP-exo analysis pipeline can be found at (https://github.com/CEGRcode/PCRPpipeline).

### Data Availability

Raw ChIP-exo sequencing files are available at NCBI GEO archive (GSE151287). Raw ChIP-seq sequencing files are available at NCBI GEO archive (GSE152144). PBM data will be available via the UniPROBE database (accession ID: LAI20A) upon publication. CUT&RUN data has been deposited at GEO: Accession GSE151326 [(5.27.20) NCBI tracking system #20907832]. Peak files for all figures are available at (https://github.com/CEGRcode/2021-Lai_PCRP)

## Acknowledgements

The Basu lab would like to thank Drs. Teresa Swayne and Emilia Laura Munteanu of the Herbert Irving Comprehensive Cancer Center Core Microscopy facility at Columbia University. The Pugh lab would like to thank Kyle Nilson for donating the peroxide stressed cells. The Shilatifard lab would like to thank Anna Whelan and Jordan Harris for technical assistance and Zibo Zhao and Marc Morgan for helpful discussions. The Bulyk lab thanks Steve S. Gisselbrecht for assistance with preparation of figures. ChIP-exo data was made available through the Cornell Institute of Biotechnology’s Epigenomic Core Facility using the Platform for Epigenome and Genomic Research (PEGR), with NIH 5R01-ES013768-12 funding for the development and dissemination of PEGR. The authors gratefully acknowledge the support of the Institute for Computational and Data Sciences at the Pennsylvania State University through an ICDS Seed Grant. This work was supported by an administrative supplement to NIAID grant 1R01AI099195 to U.B., an administrative supplement to NIH grant R21 HG009268 to M.L.B., an administrative supplement to NIH grant R01-ES013768 to B.F.P., an administrative supplement to NIH grant R01CA214035 to A.S., NIH grant R50CA211428 to E.R.S., and NIH grant R44 DE029633 to EpiCypher.

## Author Contributions

In the Basu lab, G.R. performed STORM experiments; W.Z. performed required bioinformatics analyses of STORM data; U.B. analyzed data, provided oversight and co-wrote the manuscript. In the Bulyk lab, J.T.A. and S.K.P. performed cloning, protein expression, western blots, PBM experiments, and PBM analysis; L.M. performed analysis of motif enrichment and centering; S.K.P. and L.M. prepared figures and Supplementary Tables; M.L.B. supervised research; and S.K.P., L.M., and M.L.B. co-wrote the manuscript. In the Pugh lab, TRB, KB, JM, SND, and KM performed ChIP-exo assays. PK designed and implemented the web portal. MJR and DJ performed ChIP-exo library quantitation and sequencing. WKML directed the ChIP-exo experiments, processed and analyzed the ChIP-exo, and co-wrote the manuscript writing. BFP provided oversight for ChIP-exo and co-wrote the manuscript. In the Shilatifard lab, A.P.S. conducted experiments. E.R.S. analyzed ChIP-seq data. E.R.S. and A.S. provided oversight and co-wrote the manuscript. From EpiCypher, B.J.V., K.N. and E.M. performed CUT&RUN studies, and M-C.K. provided oversight and co-wrote the manuscript.

---

**Supplementary Fig. 1.**
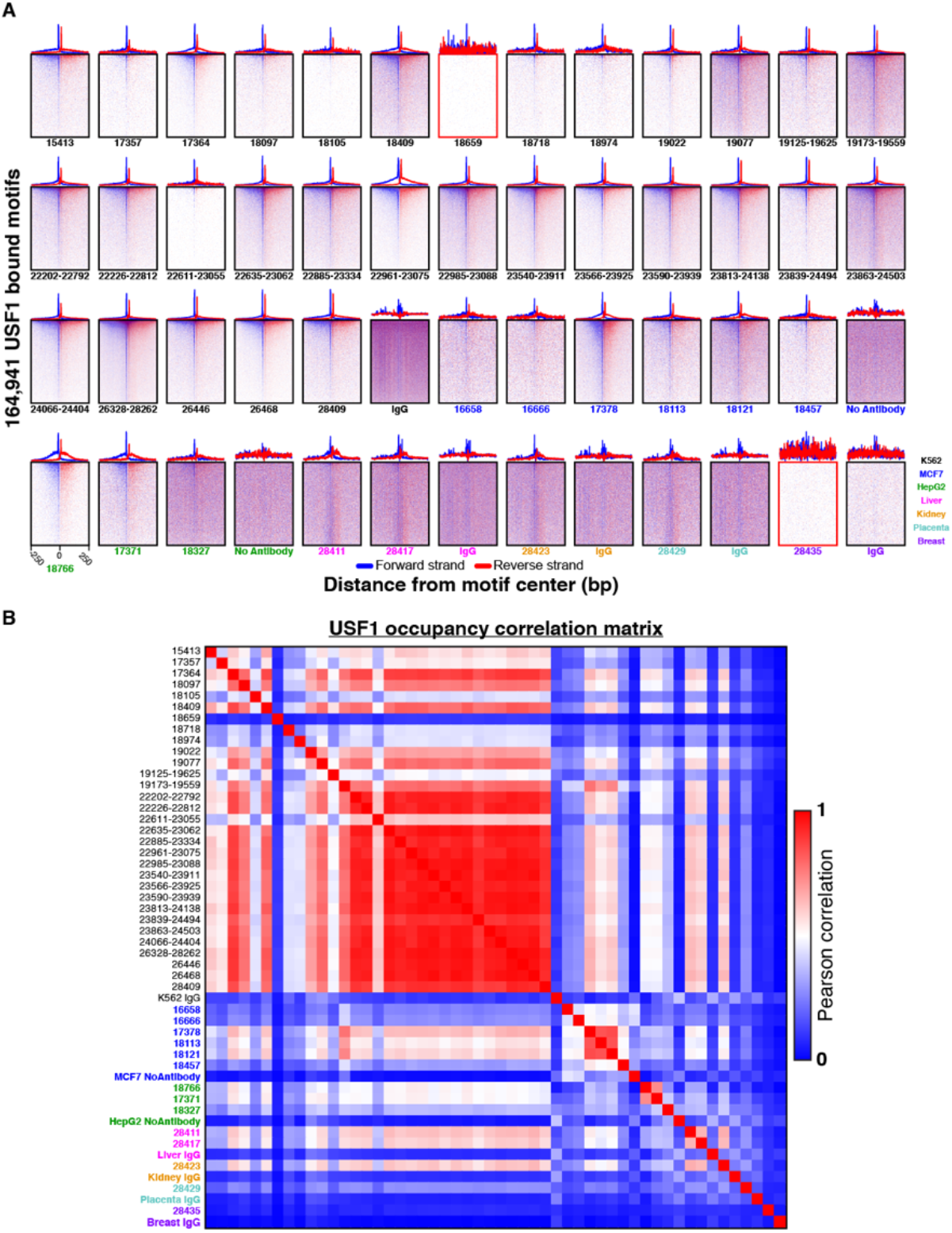
Technical robustness of the USF1 PCRP mAb in the ChIP-exo assay. (**A**) Heatmap and rowaveraged composite plots for 45 USF1 ChIP-exo experiments with the indicated project sample ID were performed in K562 (black), MCF7 (blue), HepG2 (green), human liver (pink), kidney (orange), placenta (cyan), and breast (purple). The 5’ end of aligned sequence reads for each replicate were plotted against their distance from the nearest USF1 E-box motif. These motifs were present in the union of peak-pairs across all 45 USF1 datasets for a total of 164,941 peaks that intersected with an E-box motif. Reads are strand-separated (blue = motif strand, red = opposite strand). Rows are linked across samples and sorted based on their combined rank-order average in a 500 bp bin around each motif midpoint. Matching IgG or No-Antibody control experiments for each cell type are shown. Samples 18659 and 28435 are outlined in red and represent experiments that failed to show enrichment at USF1 peaks. (**B**) Correlation matrix of USF1 technical replicates. Pearson correlation was calculated between technical replicates and negative controls using the sum of tags in a 500 bp window centered around the motif midpoint for all potential USF1 binding events. Samples are labelled and colored as defined in panel **B**.

**Supplementary Fig. 2.**
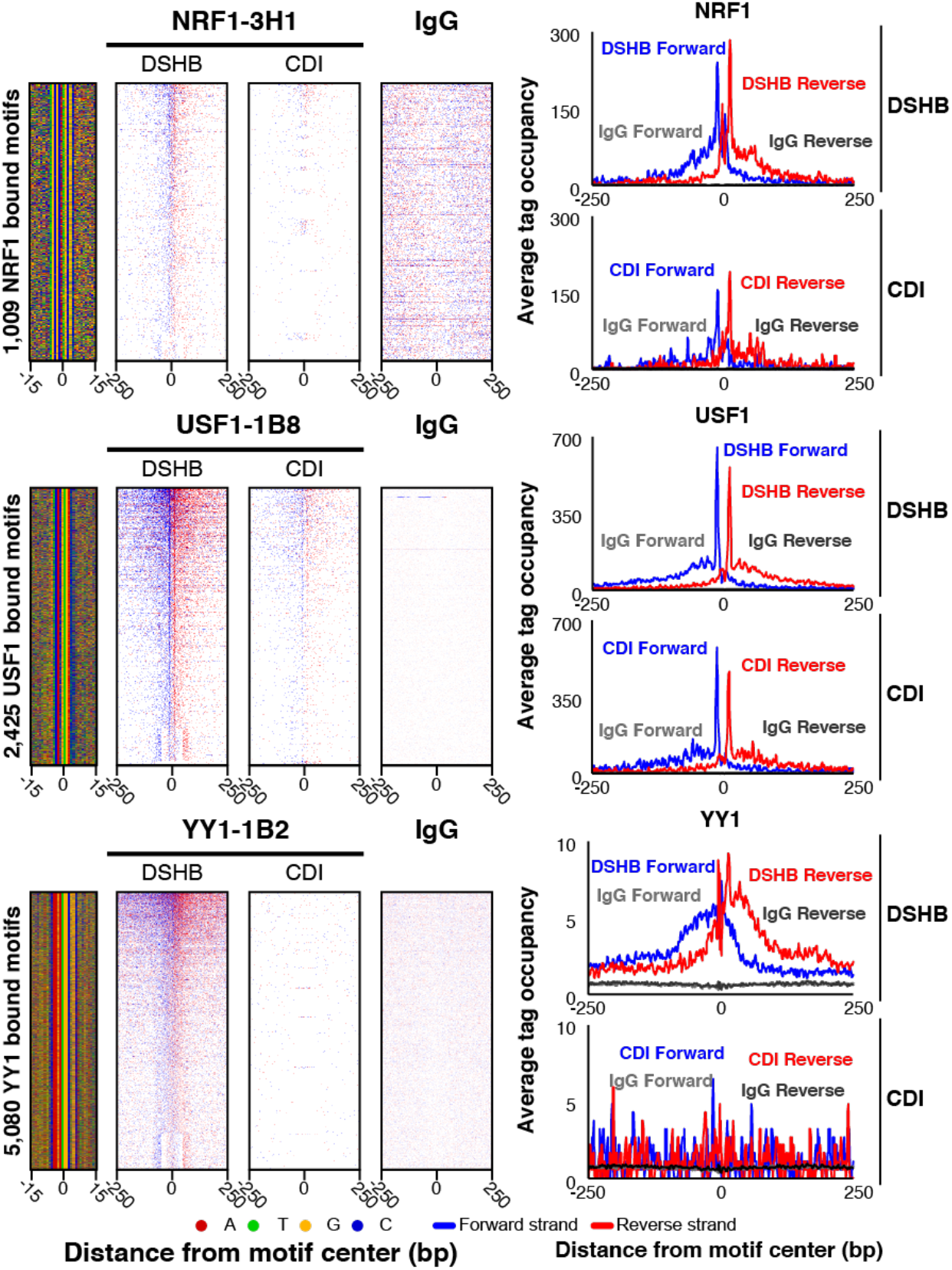
Assessment of antibody source (DSHB and CDI). DNA-sequence 4-color plots (left), heatmaps (middle), and composites (right) were generated for the indicated targets, number of bound motifs, and antibody source, tested in K562 cells. The 5’ end of aligned sequence reads for each set of experiments were plotted against distance from cognate motif, present in the union of all called peaks between the datasets for each indicated target. Reads are strand-separated (blue = motif strand, red = reverse strand). Rows are linked across samples and sorted based on their combined average in a 100 bp bin around each motif midpoint.

**Supplementary Fig. 3.**
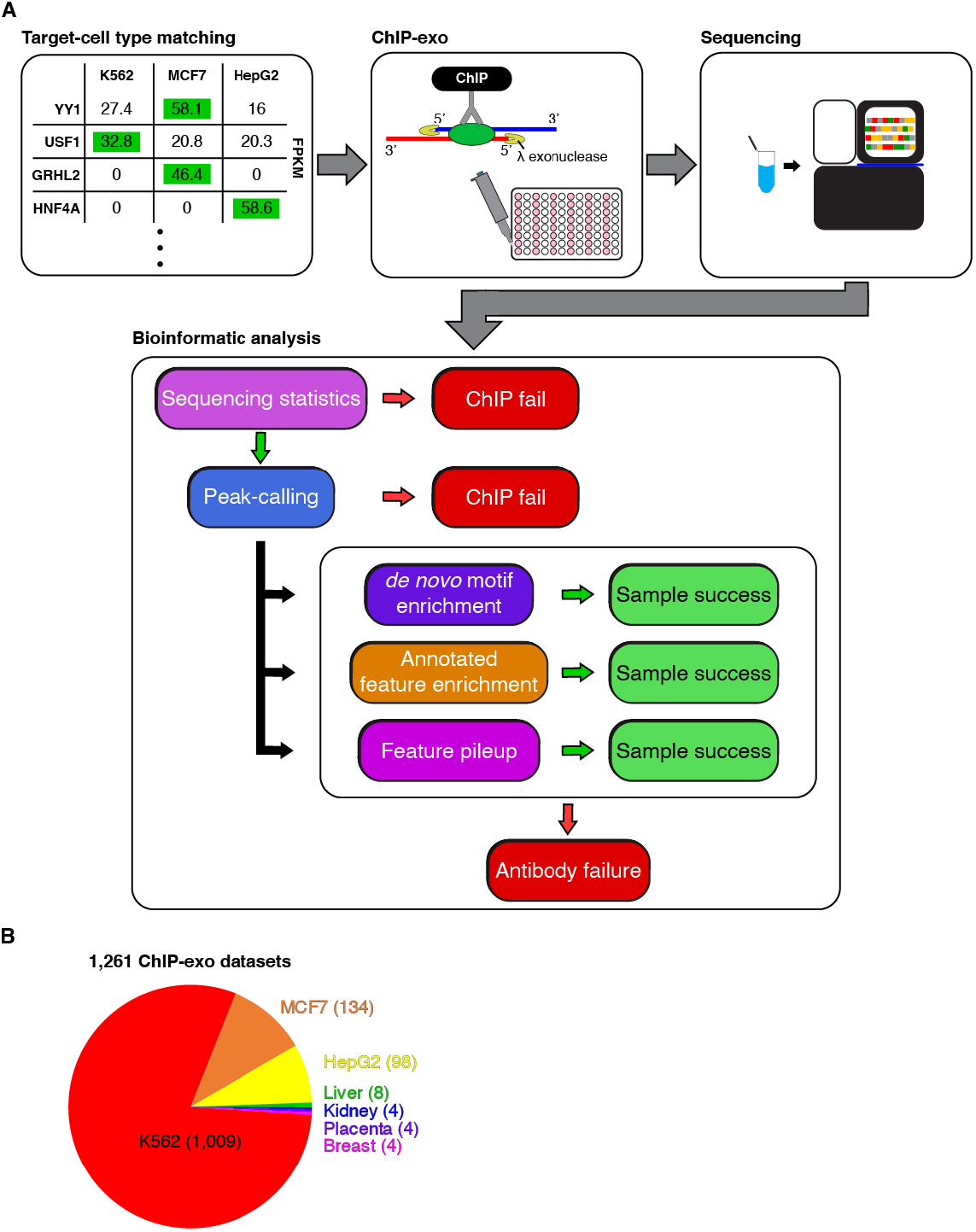
(**A**) Workflow schematic of bulk PCRP mAb testing in the ChIP-exo assay. Targets having a strong RNA-seq (Consortium 2012) expression bias towards K562, MCF7, or HepG2 were assayed in that cell line. Otherwise, they were assayed in K562. Samples were processed in cohorts of 46 plus a USF1 positive control and an IgG or “No Antibody” negative control. After high-throughput sequencing, samples were automatically processed through a bioinformatics quality control pipeline. Sample were examined for sequencing depth, library complexity (% PCR duplication), and the ability to generate significant peaks. Peaks and raw tags were then examined to identify enriched sequence motifs, localization to annotated chromatin and sequence regions, and specific enrichment at genomic features such as transcription start sites. (**B**) Pie chart shows the cell/tissue type of biological material used for 1,261 ChIP-exo datasets.

**Supplementary Fig. 4.**
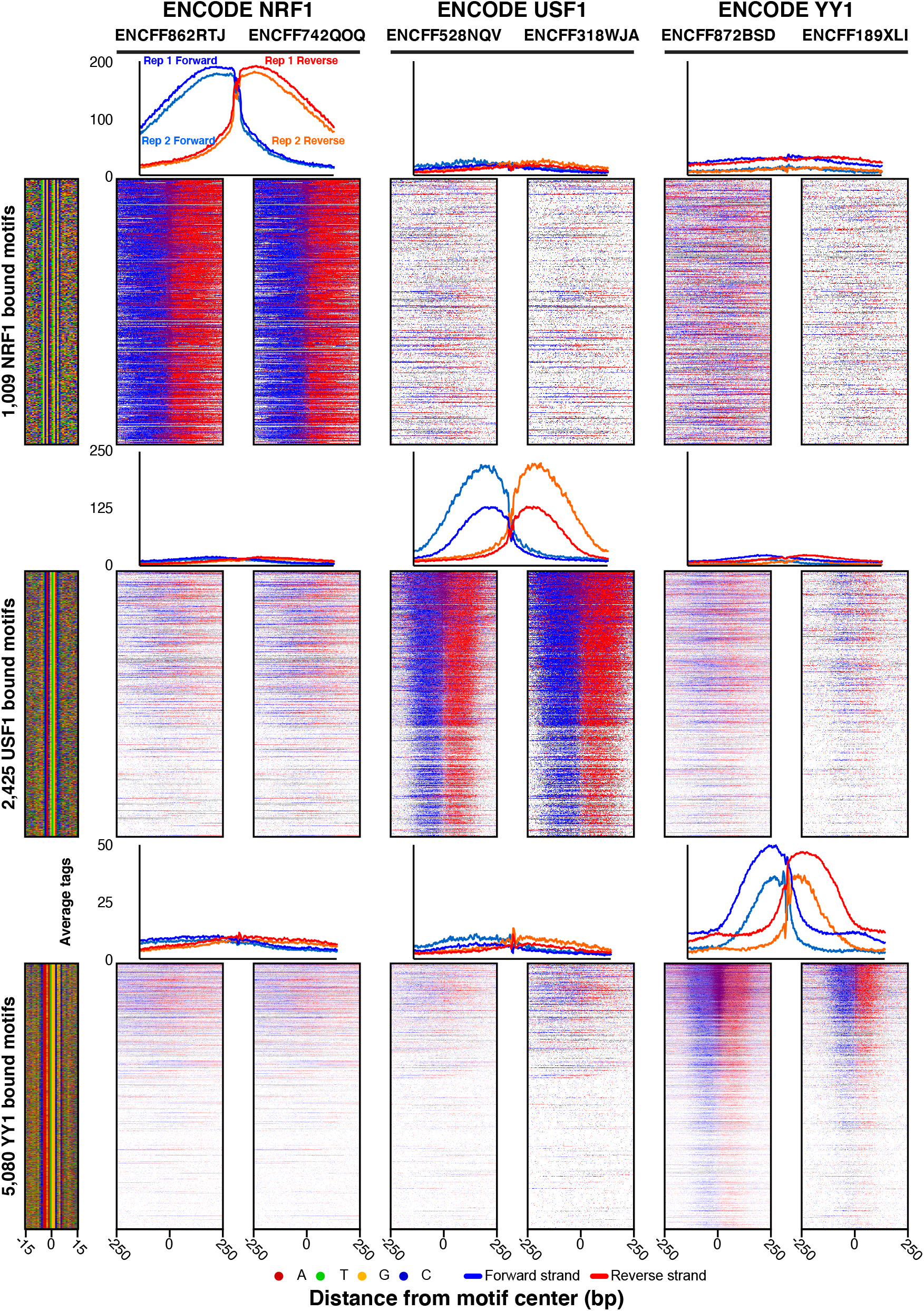
Overlay of ENCODE data at motifs defined by PCRP mAbs using ChIP-exo. ChIP-seq heatmap, composite, and DNA-sequence 4-color plots at the bound motifs defined in Figure 1 for the indicated targets in K562. The 5’ end of aligned sequence reads for each set of experiments were plotted against distance from cognate motif of target. Reads are strand-separated (blue = motif strand, red = opposite strand). Rows are linked across samples and sorted as in *Supplementary Fig. 2*.

**Supplementary Fig. 5.**
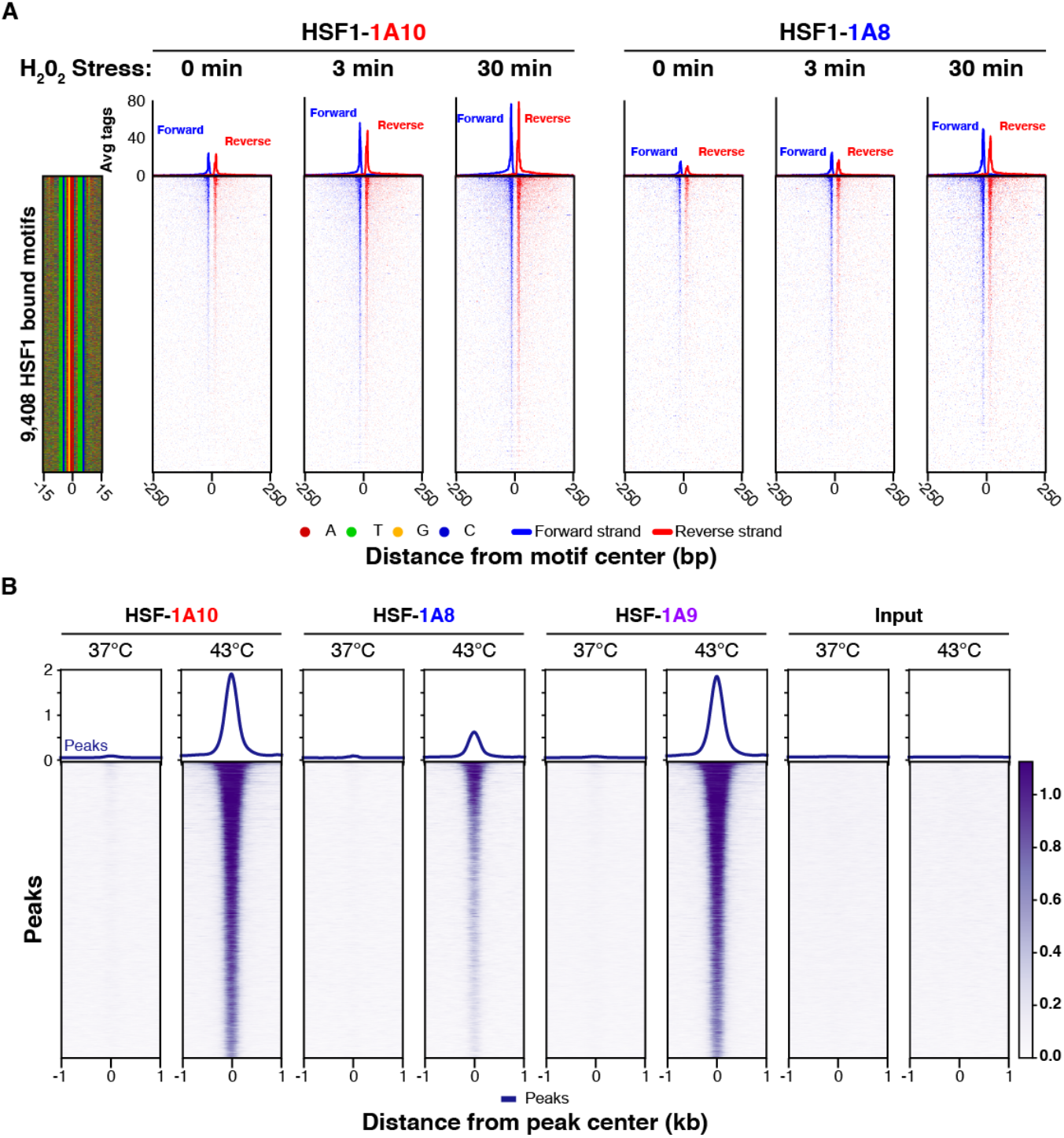
Validation of HSF1 mAb in a cell state change (stress). (**a)** ChIP-exo heatmap, composite, and DNA-sequence 4-color plots are shown for the indicated number of bound motifs for the indicated mAb in response to hydrogen peroxide treatment (0.3mM); where binding increases with treatment time) in K562 cells. The 5’ end of aligned sequence reads for each set of experiments were plotted against distance from cognate motif, present in the union of all called peaks between the datasets for each indicated target. Reads are strand-separated (blue = motif strand, red = opposite strand). Rows are linked across samples and sorted based on their combined average in a 100 bp bin around each motif midpoint. (**b)** ChIP-seq heatmap and composite plot are shown for the indicated number of bound loci for the indicated antibody hybridoma clone and input in HCT116 cells in response to 1 hr. of heat shock (42ºC) or mock (37ºC). Rows are linked across samples and sorted in descending order by mean score per region.

**Supplementary Fig. 6.**
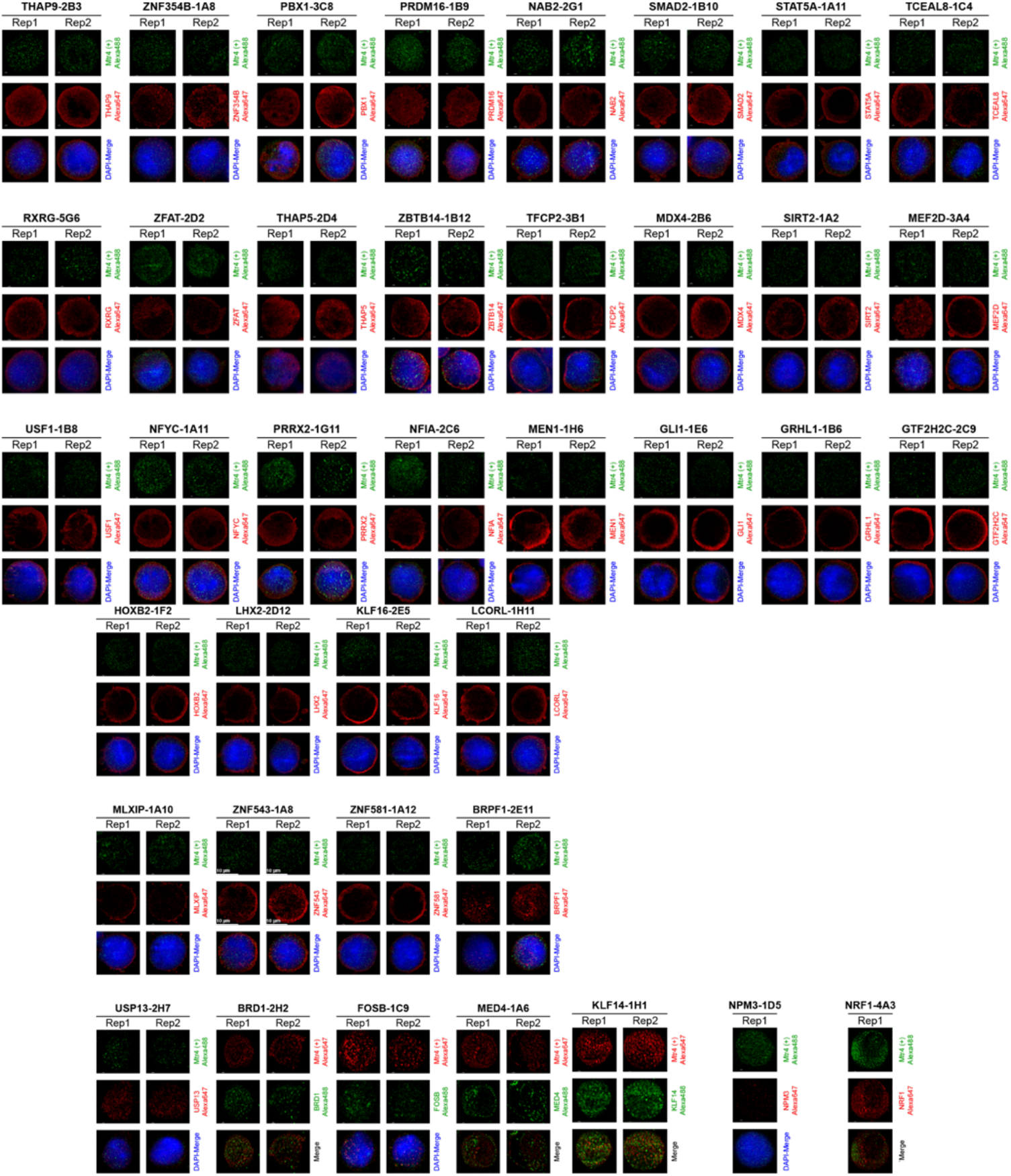
Example of STORM capabilities employing three different fluorescence channels displaying one K562 cell for nuclear localization. The STORM positive control was performed by staining the cell with commercial anti-Mtr4 (a helicase expected to be found in the nucleus and cytoplasm^37^) followed by incubation with secondary anti-rabbit conjugated to Alexa-488 (green) or Alexa-647 (red). Cells were stained with concentrated PCRP supernatant and then incubated with secondary antibody conjugated to Alexa-488 (green) or Alexa-647 (red) as labelled. DAPI nuclear staining was performed where indicated to contrast the location of the nucleus relative to sample and Mtr4 positive control staining. Note that for many of the PCRP antibodies, the antibody staining forms a ring at the periphery of the nucleus, possibly indicating non-specific binding.

**Supplementary Fig. 7.**
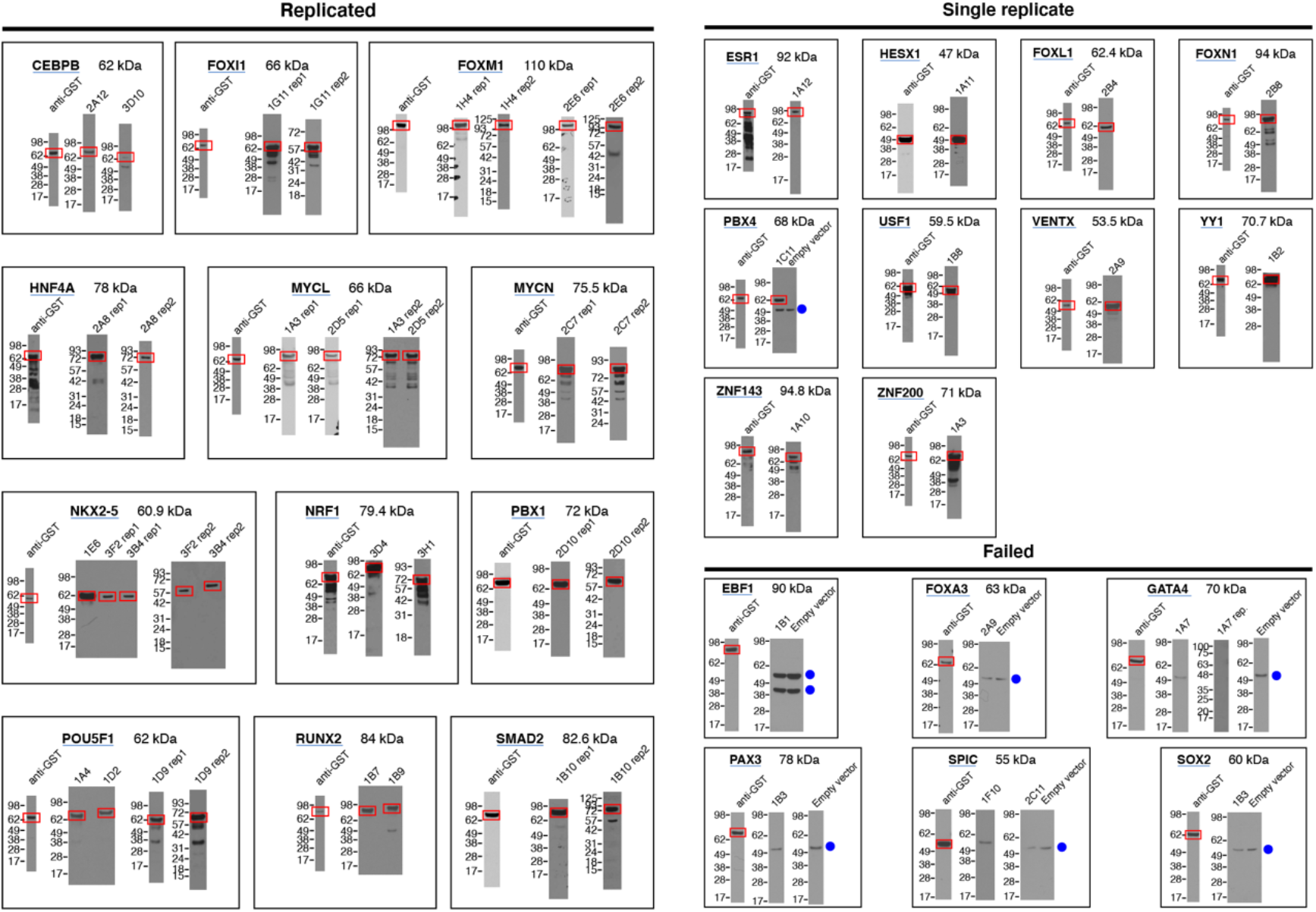
Immunoblots were conducted to assay whether PCRP antibodies can detect the full-length TF. Anti-GST immunoblots are shown for comparison. The red box indicates the correct band, corresponding to the full-length TF. An IVT negative control empty vector was assayed with PCRP antibodies and blue dot indicates cross-reactivity with the 1-Step Human Coupled IVT Kit. Replicated immunoblots (left) are composed of twelve targets that were biologically replicated with the same antibody and nine targets that were technically replicated with distinct hybridoma clones against the same target. Ten targets were assayed as a single replicate (right). The following PCRP antibodies resulted in no bands on Western blots: 1F8 (anti-FOXI1); 2C4 (anti-KLF1); 1B3 (anti-PAX3); 1C12 (anti-SMAD2); 1A7(anti-GATA4); 1A2(anti-SMAD3).

