## Supplementary figures and images for "A ChIP-exo screen of 887 PCRP transcription factor antibodies in human cells"

### Supplemental Figure 1

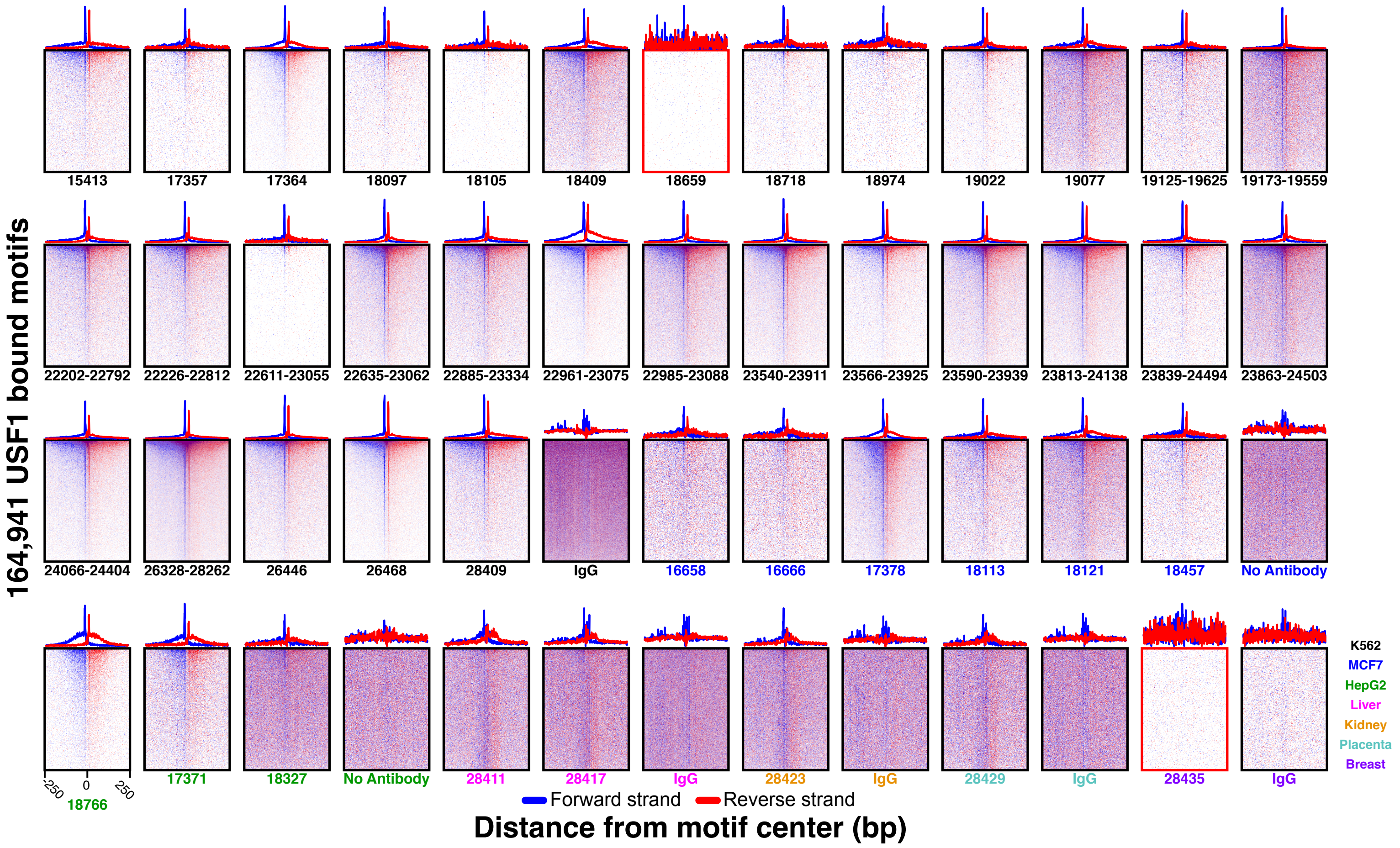

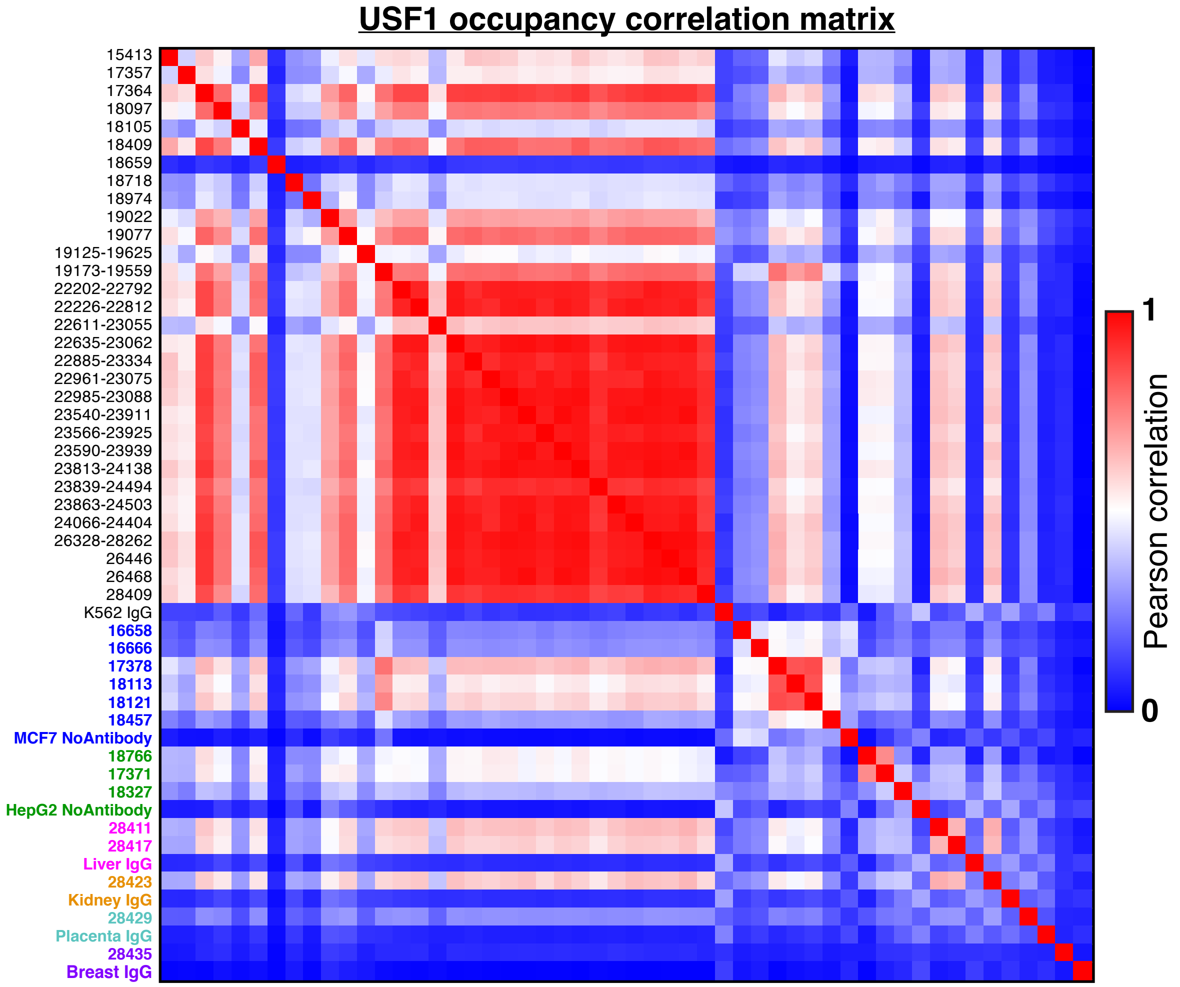

### Supplemental Figure 2

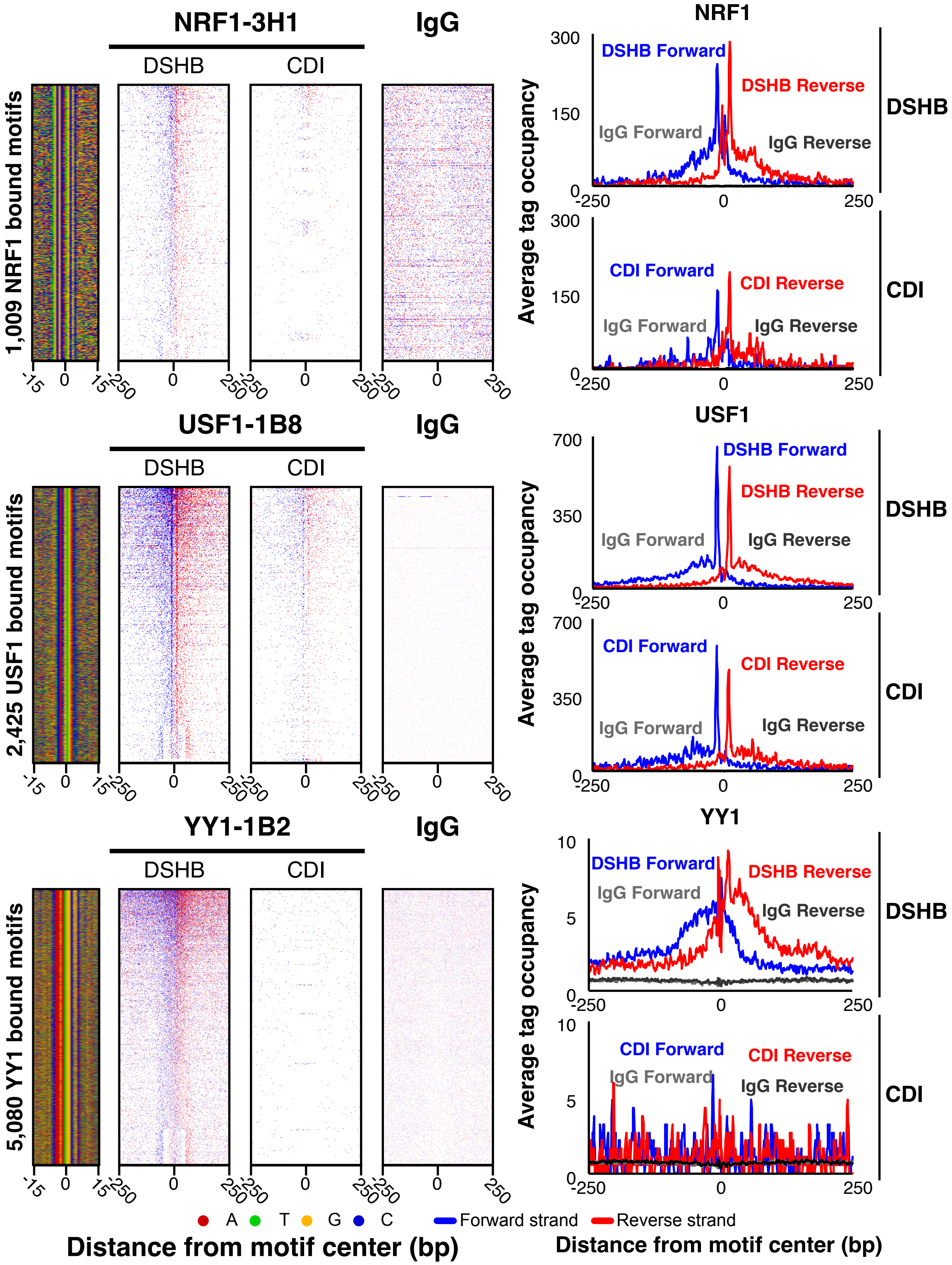

### Supplemental Figure 3

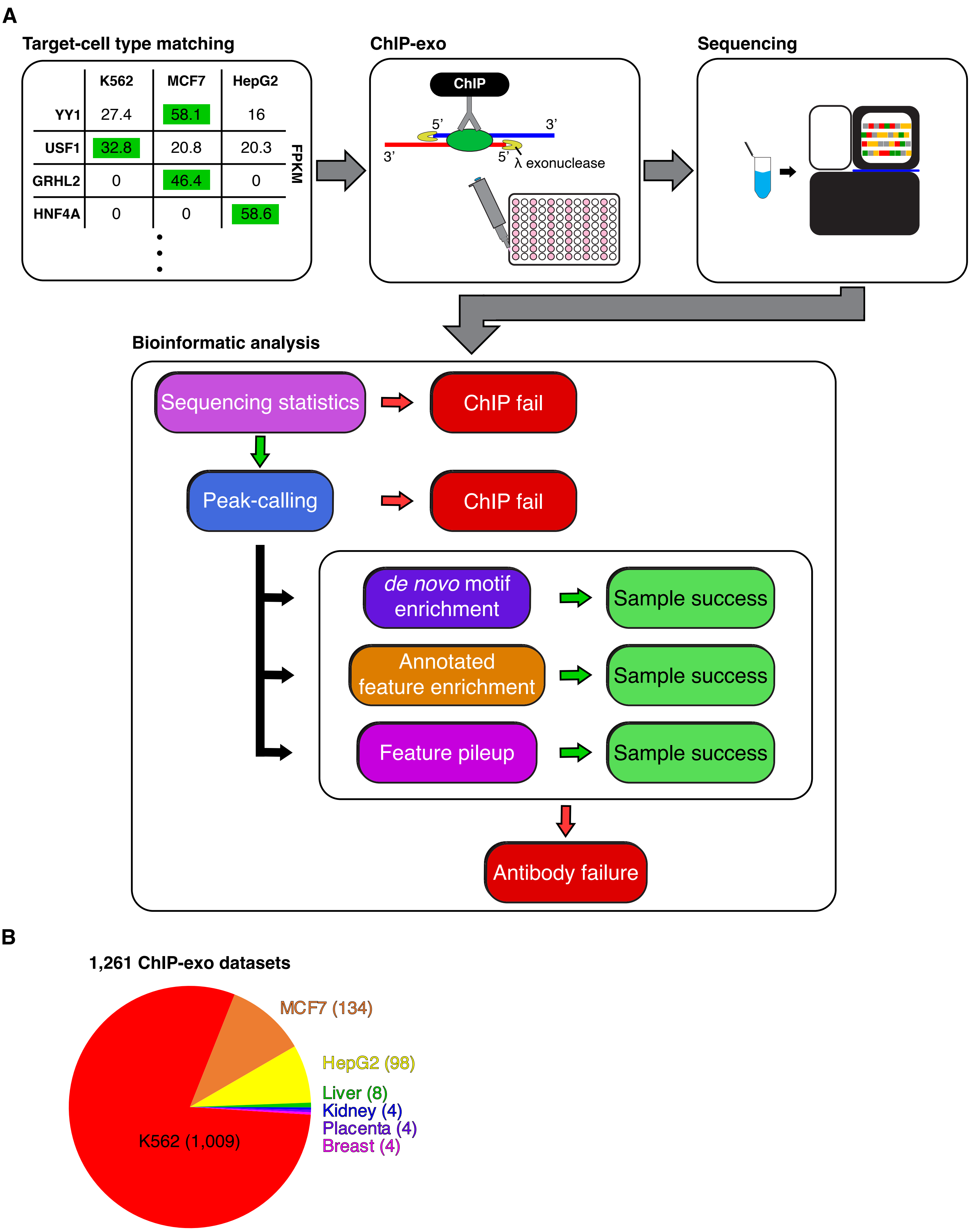

### Supplemental Figure 4

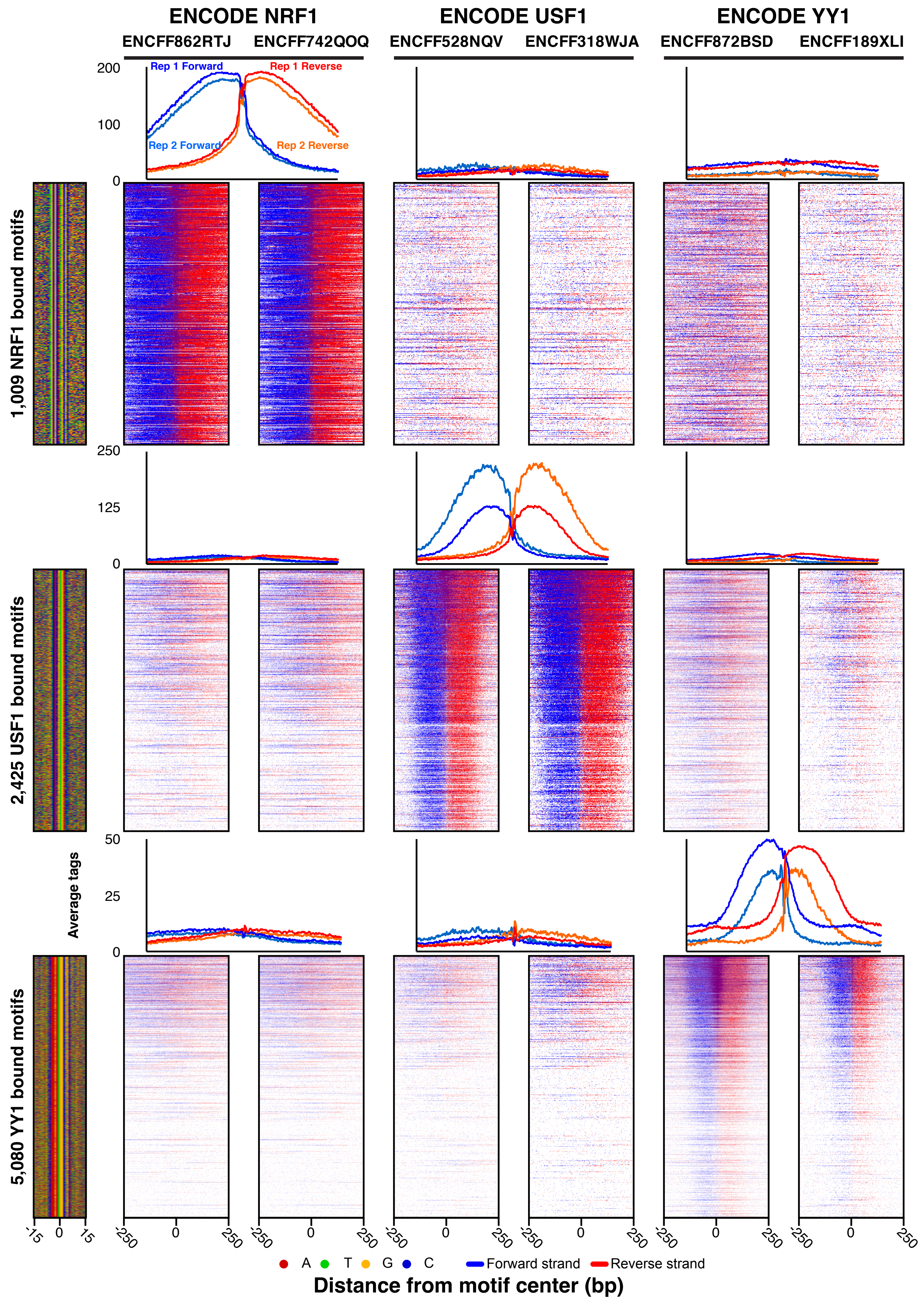

### Supplemental Figure 5

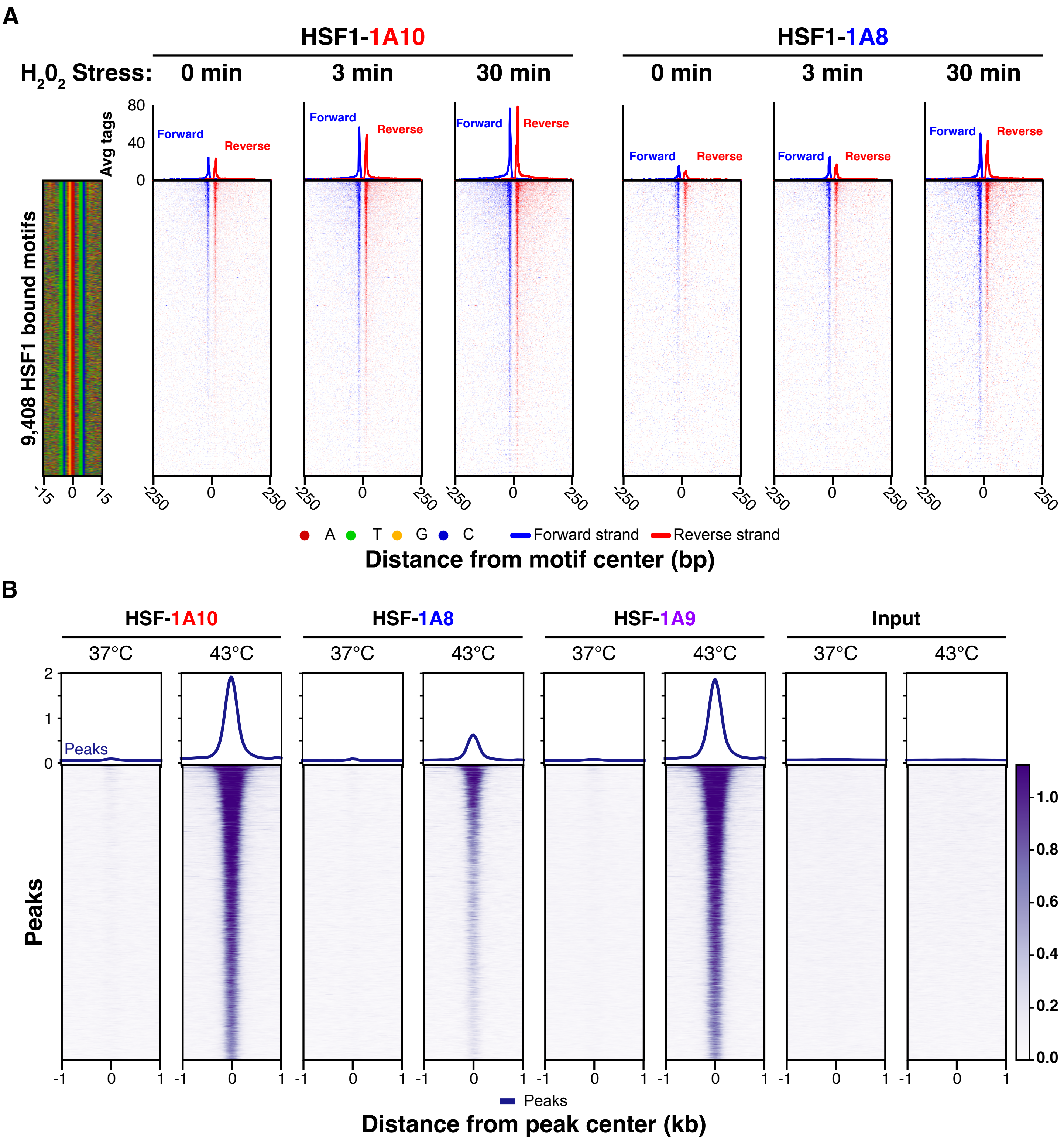

### Supplemental Figure 6

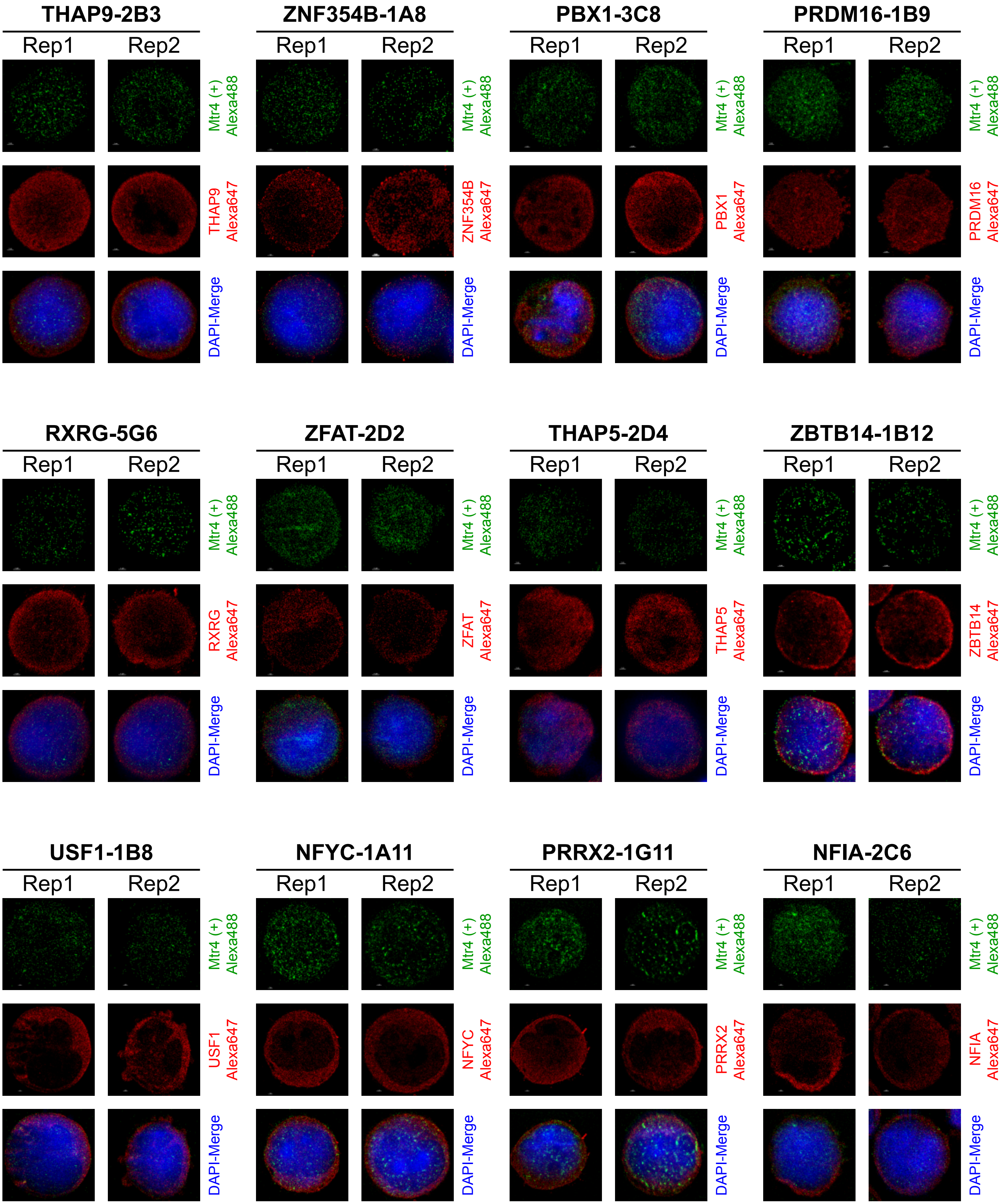

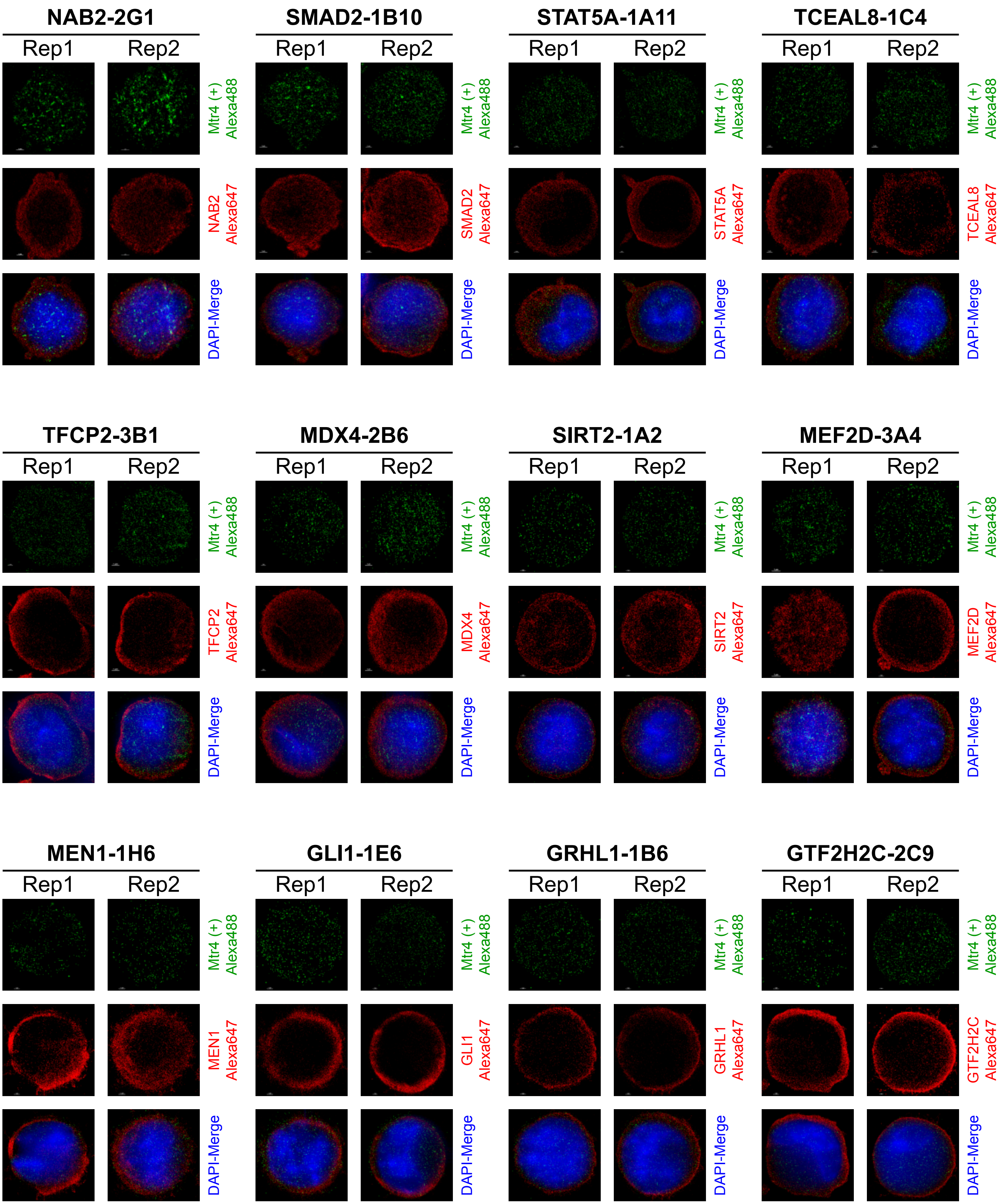

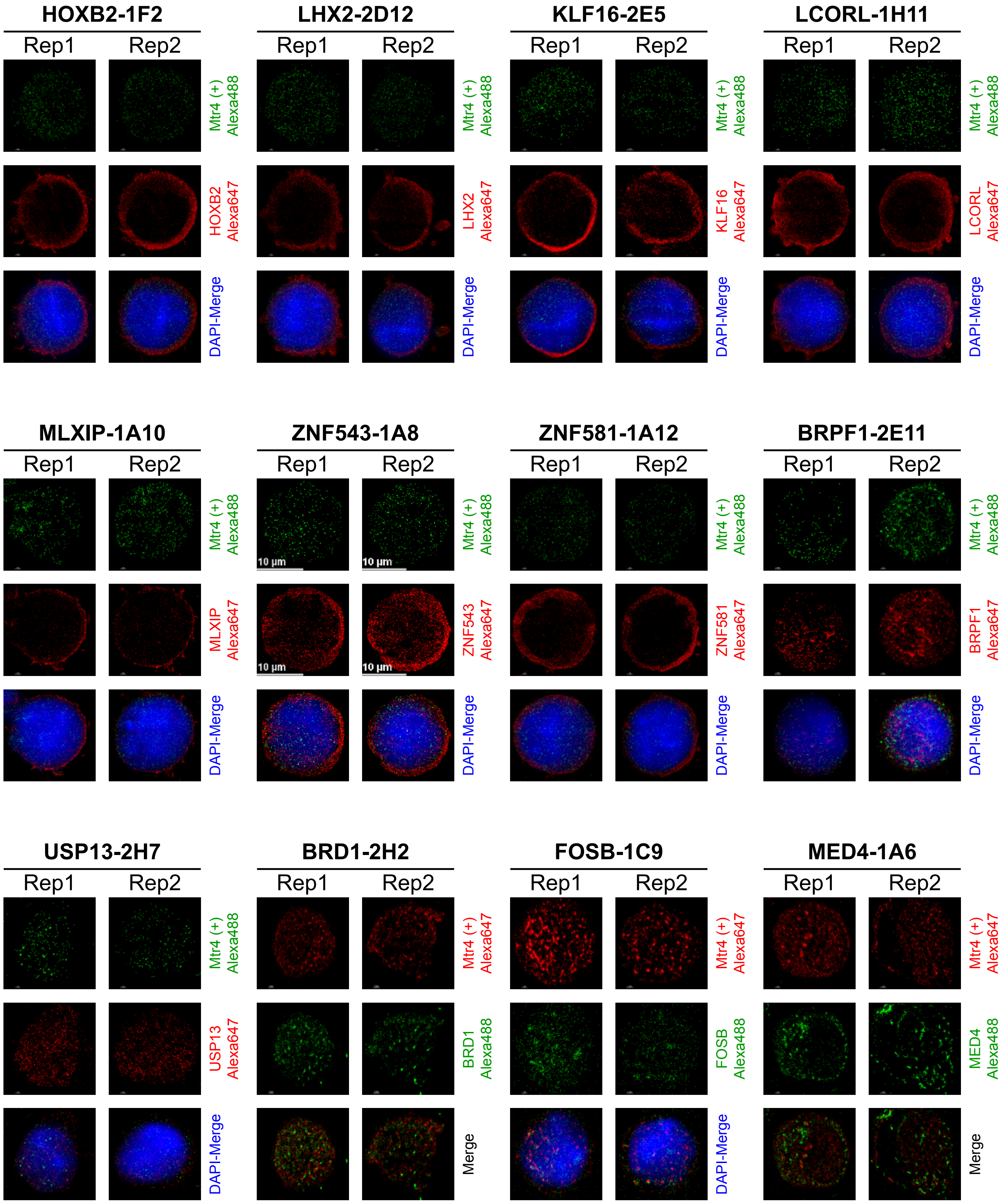

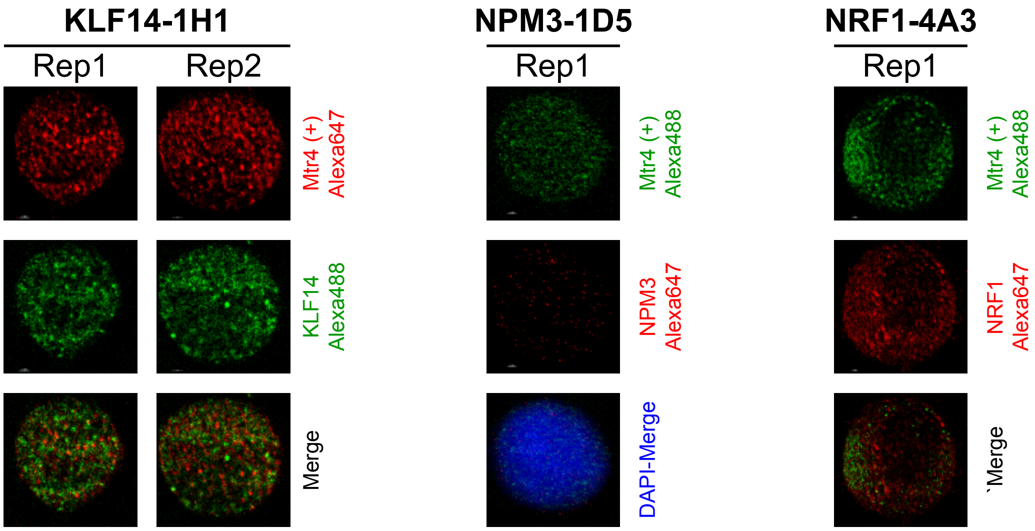

### Supplemental Figure 7

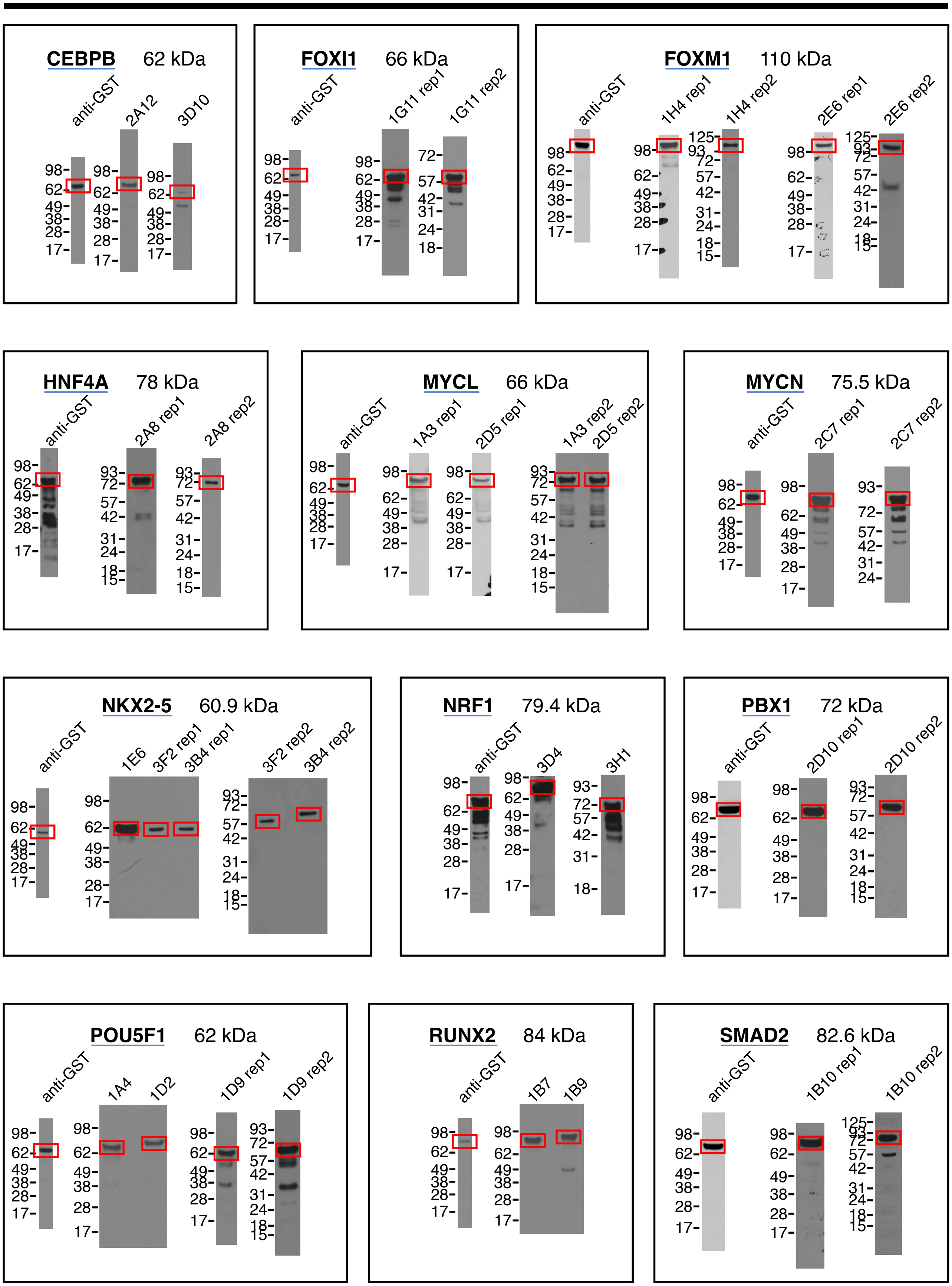

#### Single replicate

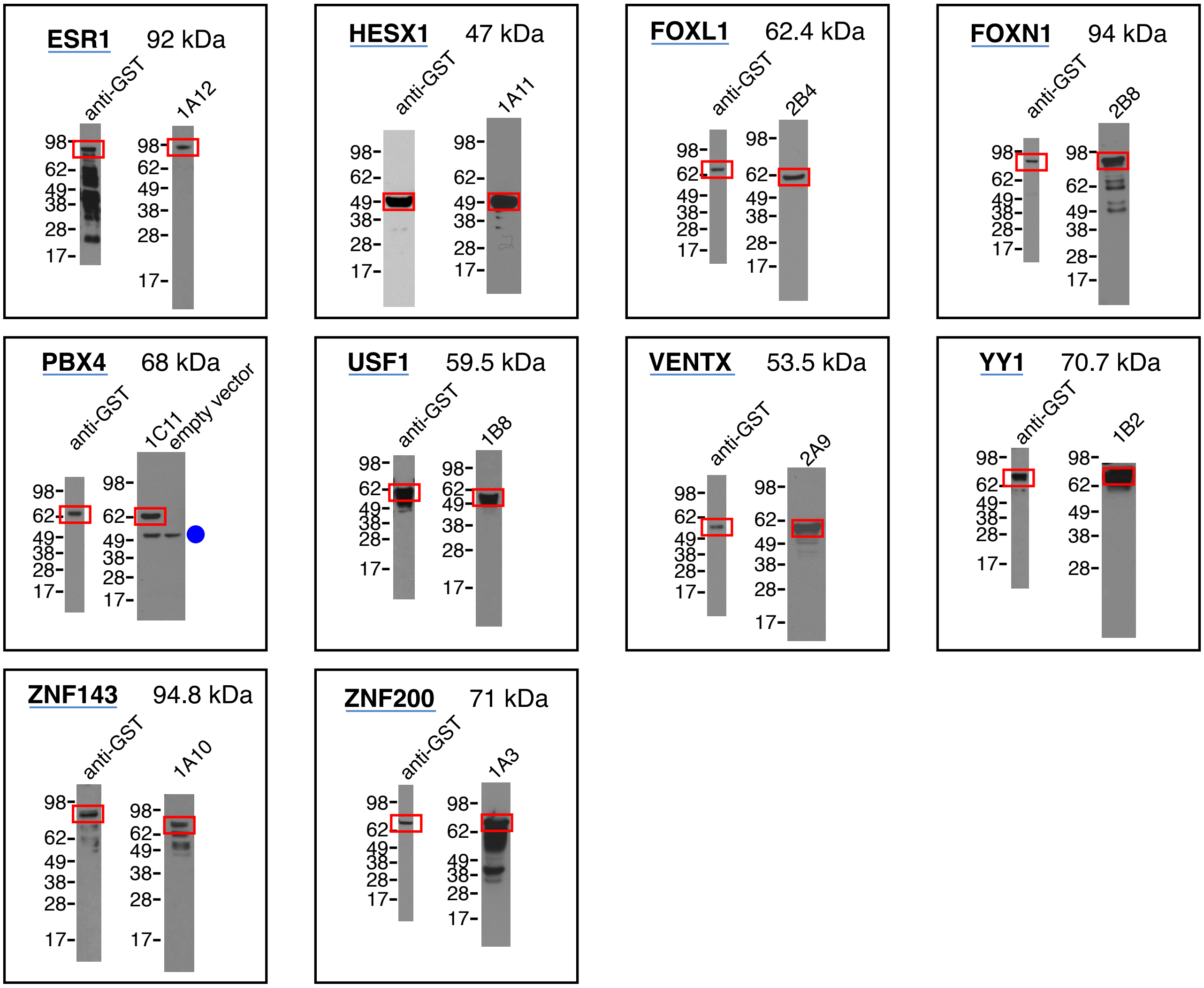


#### Failed

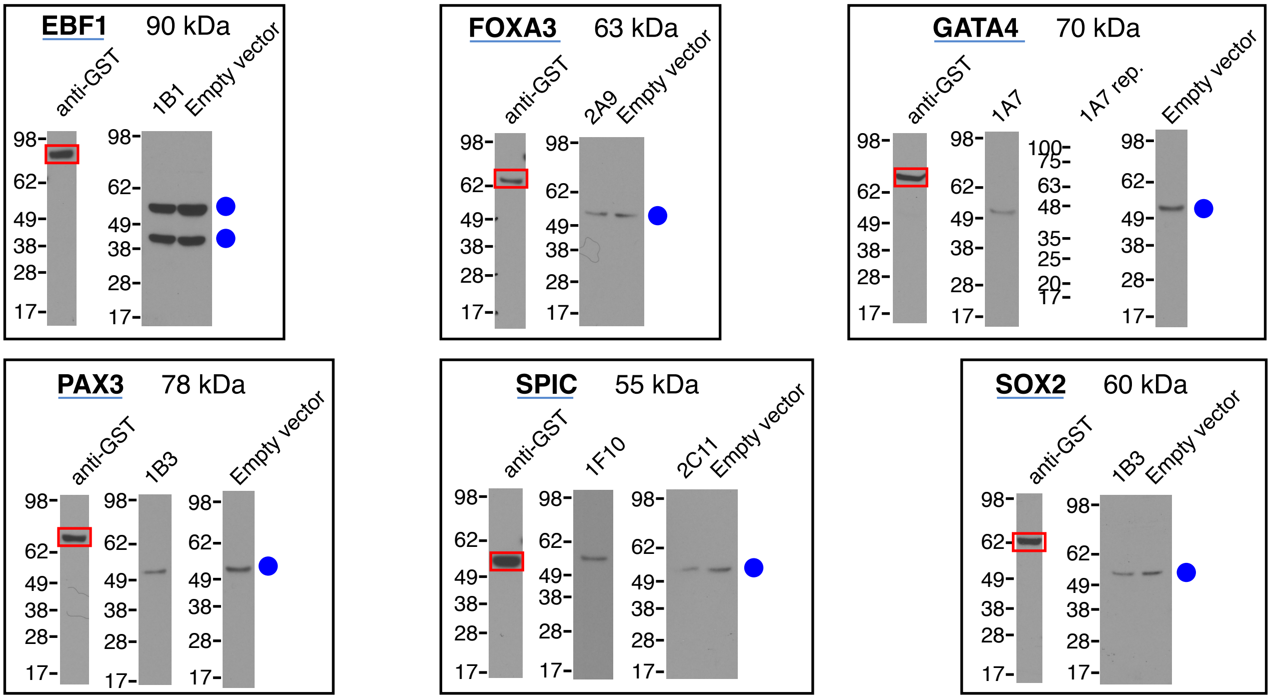
